# ADAR1 Restricts Poliovirus Infection through dual disruption of translation initiation and protein coding

**DOI:** 10.1101/2025.06.19.660622

**Authors:** Damian Kim, Frank McCarthy, Kathy Li, Robert J. Chalkley, John Pak, Joshua E. Elias, Ranen Aviner

## Abstract

Viral RNAs (vRNAs) interact with hundreds of host proteins, but how these interactions shape viral infection remains largely unknown. Here we developed a mass-spectrometry approach to identify cellular RNA-binding proteins (RBPs) that enhance or inhibit infection by modulating vRNA translation. We identified 130 RBPs that associate with polysomes in human cells infected with poliovirus, including known regulators of cap-independent translation from the viral internal ribosome entry site (IRES). We find that adenosine deaminase acting on RNA (ADAR1) is recruited to polysomes during infection and edits 3 sites in the vRNA, reducing viral replication. Two edits occur in the IRES and impair internal translation initiation, while the third edit occurs in the coding region and triggers an amino acid substitution. Incorporation of this threonine-to-alanine mutation into the viral genome attenuated replication. We conclude that ADAR1 restricts poliovirus infection by both reducing vRNA translation and introducing a recurrent coding error.

## Introduction

Cellular RNA-binding protein (RBPs) play essential roles in nearly all steps of an RNA virus life cycle, including translation, replication, processing and packaging^1–3^. Binding sites for RBPs in viral RNA (vRNA) are often under positive or purifying selection^4^, suggesting these interactions are evolutionarily conserved. A compelling example of the importance of these interactions is found in poliovirus, a positive-strand RNA (+ssRNA) virus of the *Picornaviridae* family^5^. Polio strains occupy a remarkable range of pathogenicity, from highly virulent to prophylactic. Pathogenic strains cause paralytic poliomyelitis, particularly in young children, while avirulent strains are safe and effective components of vaccines^6^. For example, the Sabin polio vaccines are based on attenuated strains that replicate efficiently in the gut, triggering an adaptive immune response, but do not infect the central nervous system or cause paralysis^7^. But what makes these strains avirulent? Evidence from cell cultures and animal models suggests that host RBPs are the key^8–13^.

All strains of polo encode a single large polyprotein of about 2,200 amino acids, which is proteolytically cleaved by viral proteases into individual monomeric viral proteins^14^. One of those, called viral protein genome-linked (VPg), acts as a primer for vRNA replication and remains covalently bound to the 5’ of genomic vRNA^14^. VPg is essential for polio replication, but it also blocks translation initiation by the canonical cap-dependent pathway^15^. Instead, the 5’ untranslated region (UTR) of polio genomic vRNA harbors an ensemble of five double-stranded RNA (dsRNA) structures that fold into an internal ribosome entry site, or IRES, that recruits ribosomes to the vRNA without requiring a 5’ cap^14^. This alternative mode of translation initiation remains active even when global cap-dependent translation is suppressed by the innate immune system^16^. However, internal translation initiation driven by picornavirus IRES is relatively inefficient and depends on multiple host RBPs, which act as noncanonical translation factors and help recruit ribosomes^10,11,17–19^.

Interestingly, the Sabin polio strains and other avirulent picornaviruses contain attenuating mutations within RBP binding sites in their IRES regions. These mutations reduce the efficiency of translation initiation, thereby decreasing viral protein production^12,13,20^. One such RBP, polypyrimidine tract-binding protein (PTB), has been implicated in the neurovirulence of polio and related picornaviruses^10,12,13,21^. While the major isoform PTBP1 enhances IRES-dependent translation, the neuron-specific isoform PTBP2 (or nPTB for neuronal PTB) does not^9^. One hypothesis posits that this difference limits viral protein synthesis in neurons infected with the Sabin strains, preventing neurological complications^9^. Although this striking example of cell type specificity may also be explained by post-translational effects on vRNA metabolism^22,23^, the idea that host RBPs function as cell type-specific barriers or bottlenecks to infection remains compelling.

Identifying host RBPs that either enhance or suppress viral protein synthesis could reveal new cellular barriers to infection and support the development of vaccines and host-directed antivirals^24,25^. In prior work, we showed that polio and other +ssRNA viruses reshape the composition of translating ribosomes (polysomes) in human cells. As infection progresses, hundreds of accessory proteins are recruited to or released from polysomes^26^. That study successfully identified molecular chaperones and modifying enzymes that promote the proper folding of viral proteins and prevent premature oligomerization and aggregation. However, it could not distinguish between polysome interactors that help viral proteins fold and those that affect translation dynamics by e.g. helping or interfering with ribosome recruitment to vRNA.

In this study, we use subcellular fractionation, RNase digestion and mass-spectrometry (MS) to discover cellular RBPs that regulate polio vRNA translation in infected human cells. We detect known enhancers and suppressors of picornavirus internal translation initiation and track their association with polysomes over the course of infection. We also identify new candidate regulators of polio vRNA translation, including adenosine deaminase acting on RNA (ADAR1). We show that ADAR1 selectively edits three major sites in the polio vRNA: two within the IRES and one in the coding region. This editing reduces polio replication by lowering translation initiation on the vRNA and introducing a detrimental coding mutation at a specific position of the viral polypeptide. ADAR1 therefore functions as a molecular barrier to polio infection by limiting the production of functional viral proteins.

## Results

### RNase digestion MS identifies host RBPs that bind polio vRNA during translation

In human cells infected with polio, a rapid and robust inhibition of cap-dependent translation is typically observed within 3 hours^26,27^ (Fig. 1A). Viral proteins, synthesized by cap-independent translation, continue to be made after global translation is inhibited (Fig. 1A). As ribosomes shift from translating host mRNAs to viral RNAs, vRNA may dominate the translatome. To confirm this, we infected Huh7 cells with type I polio (Mahoney strain) at a multiplicity of infection (MOI) of 5 particle-forming units (pfu) per cell. We then performed RNA sequencing (RNA-seq) and ribosome profiling (ribo-seq) at 2 and 4 hours post-infection (hpi)—before and after global translation shutoff. As expected, RNA fragments mapping to the vRNA genome increased from 4 to 41% of the coding pool (cytosolic mRNA+vRNA) and from 2 to 56% of the ribosome-protected pool, with host mRNAs accounting for the rest (Fig. S1A). To identify which host RBPs interact with polysomes at 4 hpi, we infected cells (MOI=5) and isolated polysomes by sucrose gradients. We digested them with RNase I, commonly used in ribo-seq analyses for its ability to degrade single stranded coding RNA, leaving ribosomes, ribosome-protected fragments and nascent polypeptide chains mostly intact^28,29^. RNase I digestion of polysomes should cleave translated vRNA in regions not protected by ribosomes, displacing RBPs that bind the vRNA without disturbing ribosome-nascent chain complexes. Digested and undigested polysomes were pelleted through sucrose cushions and their protein content was analyzed by liquid chromatography tandem MS (LC-MS/MS) (Fig. 1B and S1B).

**Fig. 1.**
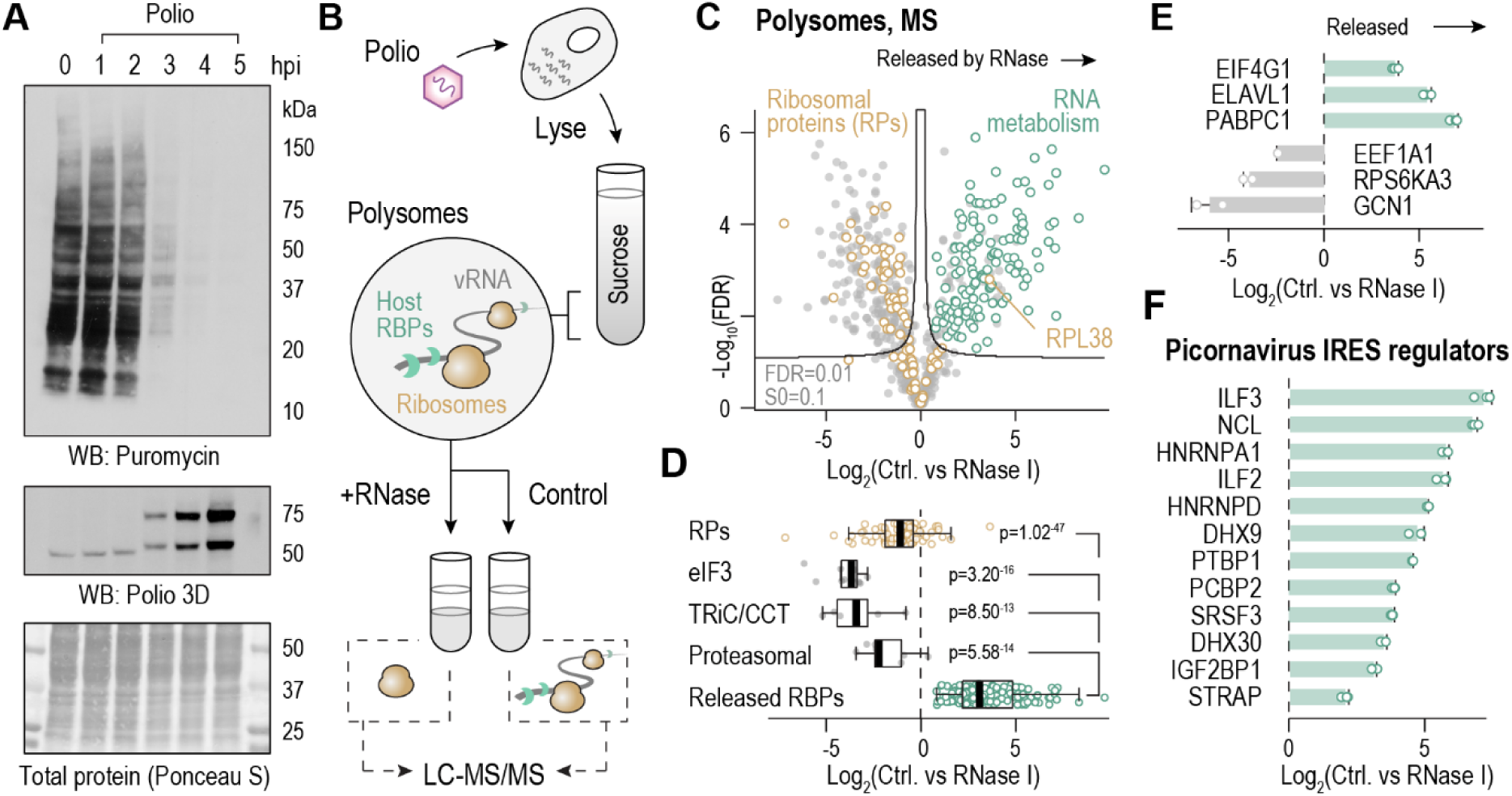
Identifying polysome-associated host RBPs during polio infection. (**A**) Huh7 cells were infected with type I poliovirus (Mahoney strain) at a multiplicity of infection (MOI)=5. Puromycin was added to tissue culture media at the indicated times (hours post-infection, hpi) to label newly synthesized proteins. Puromycylated proteins, a proxy for global translation levels, were detected by immunoblotting (n=4 independent repeats). 3D, polio nonstructural protein involved in replication (**B**) Experimental design of RNase digestion MS. Huh7 cells were infected in triplicates as in (A). Cell lysates were subjected to ultracentrifugation on sucrose gradients, and fractions containing polysomes were pooled and digested with RNase I to release RBPs. Digested and undigested polysomes were pelleted on sucrose cushions, and pellets were analyzed by liquid chromatography tandem mass-spectrometry (LC-MS/MS). (**C**) Pairwise comparisons of individual host proteins detected by MS in undigested (right) versus RNase digested (left) polysomes at 4 hpi (n=3). Proteins to the right were released from polysomes by RNase I. We consider the rest to be RNase I resistant. Yellow, cytosolic ribosomal proteins; green, proteins significantly released (FDR<0.01, S0=0.1) by RNase I that are also annotated in Gene Ontology as GOMF: *RNA binding* or GOBP: *RNA metabolic process*, excluding ribosomal proteins. (**D**) RNase sensitivity, defined as the difference in protein abundance before and after RNase treatment, for specific protein complexes associated with ribosomes. eIF3, initiation factor 3; TRiC/CCT, chaperonin; proteasomal, 26S proteasome subunits. Boxplots show the median differences in protein abundance between undigested (right) and RNase digested (left) polysomes. p, Mann-Whitney p-values. (**E-F**) Means ±SD of RNase sensitivity of proteins known to bind mRNA and vRNA during translation (E, green), proteins that directly bind the core ribosome (E, gray), and proteins known to control the activity of viral IRES in polio and related picornaviruses (F).

The most abundant protein species in both digested and undigested polysomes were cytosolic ribosomal proteins (RPs, n=78, Fig. S1C). We also detected and quantified 426 non-ribosomal proteins across all conditions and replicates (Fig. S1C and Table S1). Of those, 172 were released from polysomes by RNase I (Fig. 1C). Based on Gene Ontology (GO) annotations, 130/172 (∼75%) are implicated in RNA binding and/or RNA metabolism (Fig. 1C, green). For the purpose of this assay, we consider all other proteins to be RNase I resistant. Digestion did not displace any cytosolic RPs from polysomes, except for the large ribosomal subunit protein eL38 (RPL38) (Fig. 1C, yellow). RPL38 resides on the outer shell of the ribosome, exchanges rapidly on and off mature ribosomes, and mediates tissue-specific translation in mouse embryos^30,31^. These unique features may explain why its association with ribosomes is sensitive to RNase I, unlike all other RPs. Multi-subunit complexes that bind the core ribosome or ribosome-associated nascent chains were also RNase I resistant. These include translation initiation factor 3 (eIF3), a 13-subunit complex that remains associated with 40S subunits during translation elongation^32^ (12/13 subunits detected); and TRiC chaperonin (8/8 subunits) and 26S proteasomes (6/6 ATPase and 4/14 non-ATPase subunits), which facilitate co-translational folding and degradation of nascent chains, respectively^33,34^ (Fig. 1D). Importantly, the subset of 130 RNase-sensitive GO-annotated RBPs was well separated from ribosomal proteins and ribosome-associated protein complexes (Fig. 1C-D).

We next searched our dataset for RBPs known to bind coding RNA during translation. As expected, RNase I treatment of polysomes strongly displaced eukaryotic translation initiation factor 4G1 (EIF4G1) and poly(A) binding protein (PABPC1), which bind the 5’ and 3’ UTRs of host mRNAs^15^ and polio vRNA^35^ during translation (Fig. 1E). ELAV-like protein 1 (ELAVL1/HuR), a 3’UTR-binding protein that regulates translation of both mRNAs and vRNAs^36,37^, was also strongly displaced from polysomes by RNase I (Fig. 1E). In contrast, proteins that directly bind ribosomes during translation were resistant to RNase I. These included eukaryotic translation elongation factor 1A (EEF1A1), which delivers incoming amino acids^38^; ribosomal protein S6 kinase (RPS6KA3) that controls translation initiation^39^; and stalled ribosome sensor GCN1, which binds stalled ribosomes^40^ (Fig. 1E, gray). Taken together, RNase I digestion of polysomes from polio infected cells released host RBPs without displacing major components of the ribosome-nascent chain complex.

We then searched our dataset for RBPs known to bind and modify IRES function in polio and related picornaviruses. Proteins displaced from polysomes by RNase I included known enhancers of IRES-mediated vRNA translation e.g. PTBP1^12^, PCBP2^41^, NCL^42^, HNRNPA1^43^, DHX9^44^, SRSF3 (SRp20)^45^, IGF2BP1^46^ and STRAP (Unr-IP)^47^ (Fig. 1F). HNRNPD (AUF1), which suppresses IRES-mediated translation in polio^48^; and ILF2/3 (NF45/NF90), which suppress IRES-mediated translation in a related picornavirus, human rhinovirus type 2 (HRV2)^49^, were also displaced from polysomes by RNase I (Fig. 1F). Thus, IRES trans-acting factors (ITAFs) associate with polysomes in polio infected cells and can be detected using RNase digestion MS.

### Temporal dynamics of polysomal RBPs during polio infection

During infection, increased translation of vRNA (Fig. 1A, S1A) may lead to increased polysome association of some RBPs. To determine which of the RBPs we identified using RNase I also change in polysome abundance during infection, we re-analyzed our previously published MS dataset from polio infected Huh7 cells^26^. We used this dataset, generated under similar conditions, to compare the composition of undigested polysomes in uninfected cells and at 2, 3, and 4 hpi. Consistent with a shutoff of cap-dependent translation, infection reduced the levels of cytosolic RPs—and therefore, translating ribosomes—in the polysome pool (Fig. 2A, and Fig. 2B, top left, yellow). This reduction was observed at 3 and 4 but not 2 hpi, coinciding with translation shutoff (Fig. 1A). Of the RBPs identified in Fig. 1C, 17, 28 and 51 were recruited to polysomes at 2, 3 and 4 hpi, respectively (Fig. 2A, green). This subset included known ITAFs^50^, both enhancers (PTBP1, PCBP1, STRAP, SRSF3, SRSF7/9G8 and HNRNPC) and suppressors (HNRNPD and ILF2/ILF3) (Fig. S2A).

**Fig. 2.**
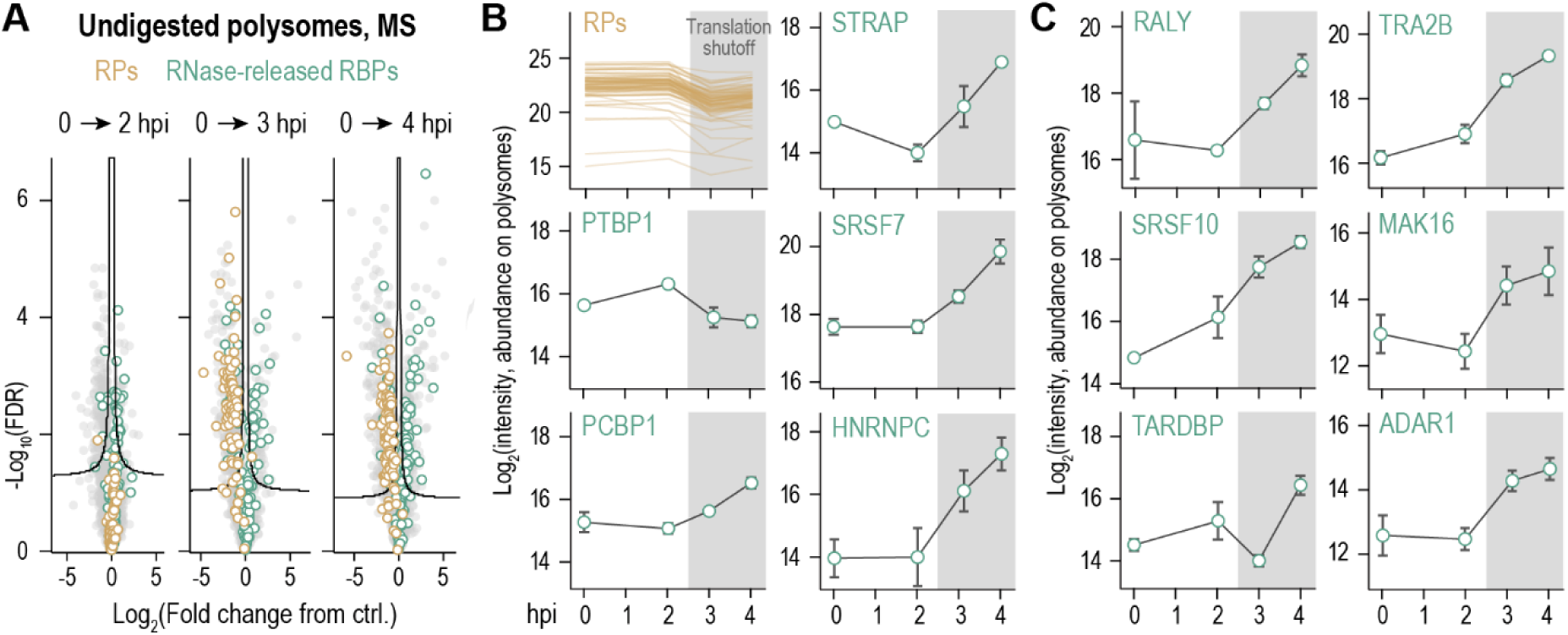
Temporal dynamics of polysomal RBPs during polio infection. (**A**) Pairwise comparisons of individual host proteins quantified by MS in undigested polysomes at the indicated times of infection (right side of each plot) compared to uninfected controls (left side) (n=3 each). (**B-C**) Time course of polysome association for ribosomal proteins (RPs, yellow) and RNase-released RBPs (green). Shown are the means of all cytosolic ribosomal proteins (RPs, yellow) or means ±SD (green) at each timepoint (n=3 each). Shading reflects global translation shutoff between 2 and 3 hpi. B, known polio ITAFs; C, new potential regulators of polio IRES identified in this analysis.

Interestingly, these ITAFs showed different temporal patterns of polysome association (Fig. 2B and S2B). For example, while PTBP1 was mildly recruited at 2 hpi and then partly displaced, PCBP1 was only recruited at 3 hpi and remained high at 4 hpi (Fig. 2B). Others followed more complex patterns of recruitment and depletion, including HNRNPA1, HNRNPA2B1 and SRSF3^51^ (Fig. S2A), pointing to a complex relationship between polysomal vRNA and the RBP it binds during infection. Many ITAFs are otherwise involved in pre-mRNA processing, particularly alternative splicing^52^. Our analyses identified new splicing-related RBPs that were both recruited to polysomes during infection and released from polysomes by RNase I. It is tempting to speculate that these, including RALY, SRSF10, TARDBP, TRA2B and MAK16 (Fig. 2C), could represent new regulators of polio vRNA translation.

### ADAR1 is recruited to translating polysomes during polio infection

ADAR1 (Fig. 2C and S2A), one of the RBPs most strongly released from polysome by RNase I (>100 fold, Table S1), was also enriched on polysomes at 3 and 4 hpi relative to uninfected controls (Fig. 2C). ADAR1 functions in multiple steps of RNA metabolism, including splicing^53^. ADARs are so-called “RNA editors” that convert adenosines to inosines in coding and noncoding RNAs and play important roles in innate immunity and viral infection^54–56^. The sequence specificity of ADARs is poorly understood, but they all recognize and edit adenosines in or around RNA duplexes. Polio vRNA genome forms multiple intramolecular duplexes that could act as potential ADAR1 substrates, located in both the 5’UTR and coding region^57^. Indeed, ADAR1 was found to interact with the structured 5’UTR of HRV2 in an RNA interactome analysis^44^.

ADAR1 exists as a short constitutive isoform (p110) and an N-terminal extended, interferon stimulated isoform (p150)^54^. Based on peptide-level MS data, we find that both ADAR1 isoforms bind polysomes in polio infected cells (Fig. 3A). Still, peptides mapping to the unique N-terminal tail of p150 were less abundant than most of those mapping to the common region shared by both isoforms (Fig. S3A). Furthermore, neither isoform was induced nor depleted within 6 hours of infection (Fig. S3B). Thus, either of the two isoforms could account for the increase in polysomal ADAR1. To determine which isoform was localized to polysomes, we infected Huh7 cells (MOI=5), harvested them at 3 hpi, fractionated cell lysates on sucrose gradients and analyzed rRNA and protein content in each fraction by UV absorbance and immunoblotting, respectively. Translation shutoff reduced the overall abundance of polysomes at 3 hpi (Fig. 3A) and shifted ribosomal protein P1 (RPLP1) from the heavy polysome to 80S fractions of the gradients (Fig. 3B-C). In contrast, the majority of ADAR1 p110 did not co-sediment with RPLP1 in uninfected cells but shifted deeper into the gradients upon infection (Fig. 3B-C). p150 showed a similar shift but was overall much less abundant than p110 (Fig. S3C). Another ADAR member, ADARB1 or ADAR2, was also recruited to polysomes during infection, at levels 5-fold lower than ADAR1 (Fig. S3D). Therefore, multiple ADAR proteins may be involved in polio infection. As a control for polysome association, we also tested the distribution of TRMT6/TRMT61A, a two-subunit protein complex that remained associated with but not recruited to polysomes during infection (Fig. S3C, bottom; and Table S1). Unlike ADAR1, both members of the TRMT6/TRMT61A complex shifted to the lighter fractions of the gradients upon polio infection (Fig. 3B).

**Fig. 3.**
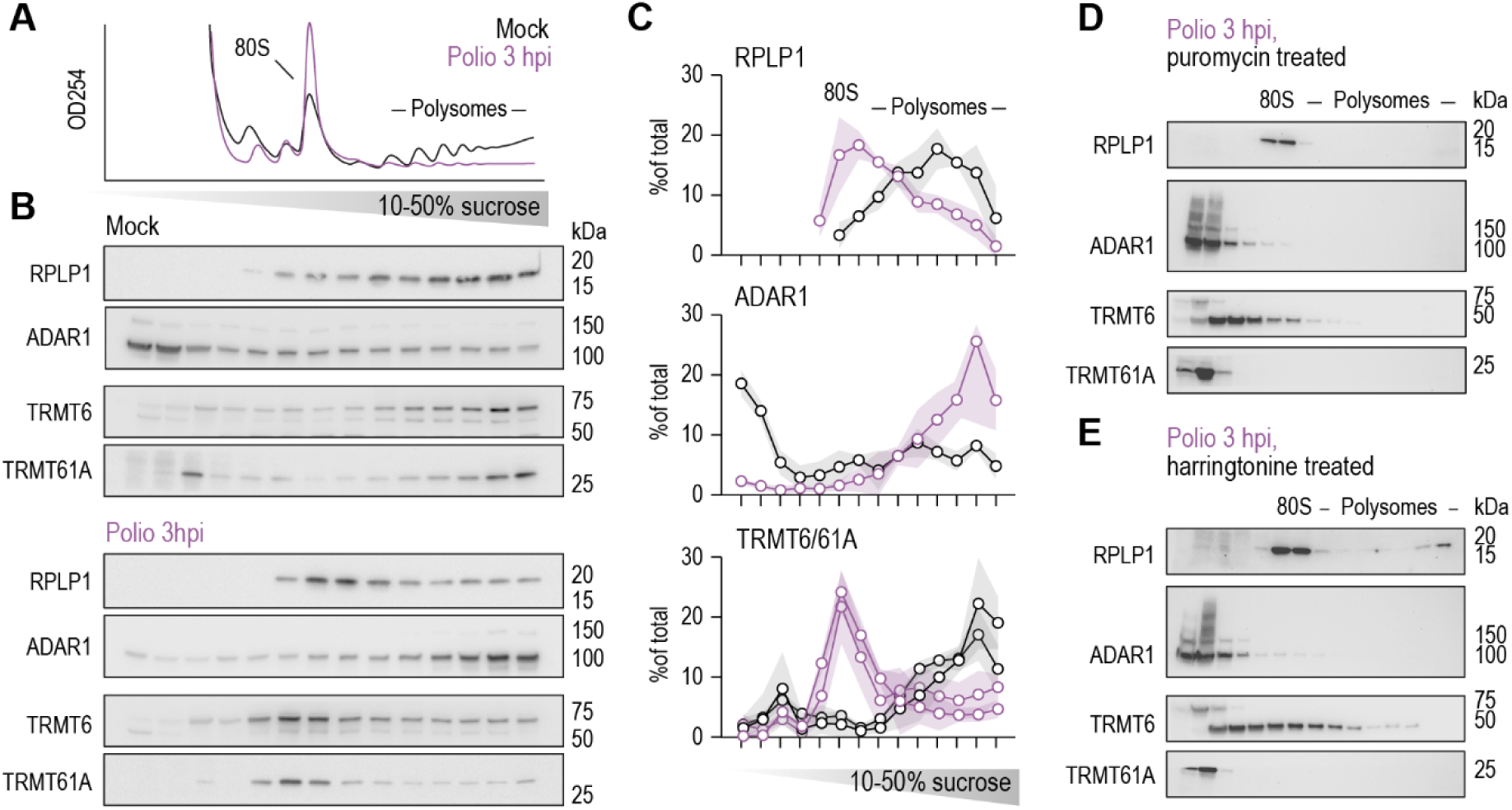
ADAR1 binds translating polysomes during polio infection. (**A**) Huh7 cells were infected (MOI=5) and lysates were fractionated on 10-50% sucrose gradients with constant monitoring of rRNA absorbance. (**B**) Gradient fractions were probed by immunoblotting for the indicated proteins. RPLP1, 60S ribosomal protein P1. Blots are representative of n=3. (**C**) Summary densitometry from (B). Percent of total was calculated by dividing the density value in each fraction by the summed densities of all fractions. Means in circles, interquartile range in shade. Densitometry was performed separately for TRMT6 and TRMT61A, and the plots were overlaid. (**D-E**) Huh7 cells infected as above were treated with 10 μg/mL puromycin (D) or 2 μg/mL harringtonine (E) for 15 min to induce termination of elongating ribosomes and deplete polysomes. Blots are representative of n=2 each.

Next, we infected cells as above and pretreated them with either puromycin or harringtonine before harvesting. These drugs trigger termination and runoff of elongating ribosomes, respectively. If ADAR1 binds active polysomes, these treatments should displace it by disassembling preexisting polysomes and preventing the formation of new ones. Both puromycin and harringtonine shifted RPLP1 to the 80S peak, consistent with induced translation termination (Fig. 3D-E). Both treatments also shifted ADAR1 to the lightest fractions of the gradients, where it no longer co-sedimented with RPLP1 (Fig. 3D-E). TRMT6/TRMT61A also shifted in response to puromycin and harringtonine (Fig. 3D-E), confirming these proteins still bind translating polysomes during infection (Fig. S3C). Taken together, this suggests that ADAR1—and particularly p110—binds co-translationally to polysomes in polio infected cells.

### ADAR1 edits specific sites on polio vRNA

To determine whether polio vRNA is edited by ADAR1, we first measured its inosine content. We infected cells (MOI=5), extracted total RNA and used beads coated with an anti-inosine antibody to affinity purify any RNA that contains inosines. Bead-bound vRNAs were quantified by qPCR using primers that amplify regions in either the 5’ UTR or coding sequence (CDS). If viral genomes are edited by ADAR1 during infection, they should be enriched in inosine IPs. Indeed, viral genomes were enriched several orders of magnitude in inosine pulldowns compared to IgG controls at both 3 and 4 hpi (Fig. 4A). Knockout (KO) of ADAR1 reduced but did not eliminate capture of viral genomes by anti-inosine beads (Fig. 4B), suggesting they may still be edited at lower levels. This could be due to residual ADAR1 protein in KO cells or redundancy with other polysome-recruited ADARs e.g. ADARB1 (Fig. S3C). Regardless, lower levels of inosine-containing vRNA in ADAR1_KO_ cells could also be explained by lower viral replication in the absence of ADAR1. To test this, we infected ADAR1_WT_ and ADAR1_KO_ cells (MOI=0.1) and measured intracellular vRNA levels, relative to GAPDH, by qPCR. ADAR1 KO decreased the levels of inosine-containing viral genomes by about 4 fold (Fig. 4B) but increased the overall levels of viral genomes by up to 3 fold (Fig. 4C). The same was observed in a second ADAR KO line, generated by another CRISPR guide (Fig. S4A). Therefore, we conclude that ADAR1 edits adenosines into inosines on polio vRNA and restricts viral replication.

**Fig. 4.**
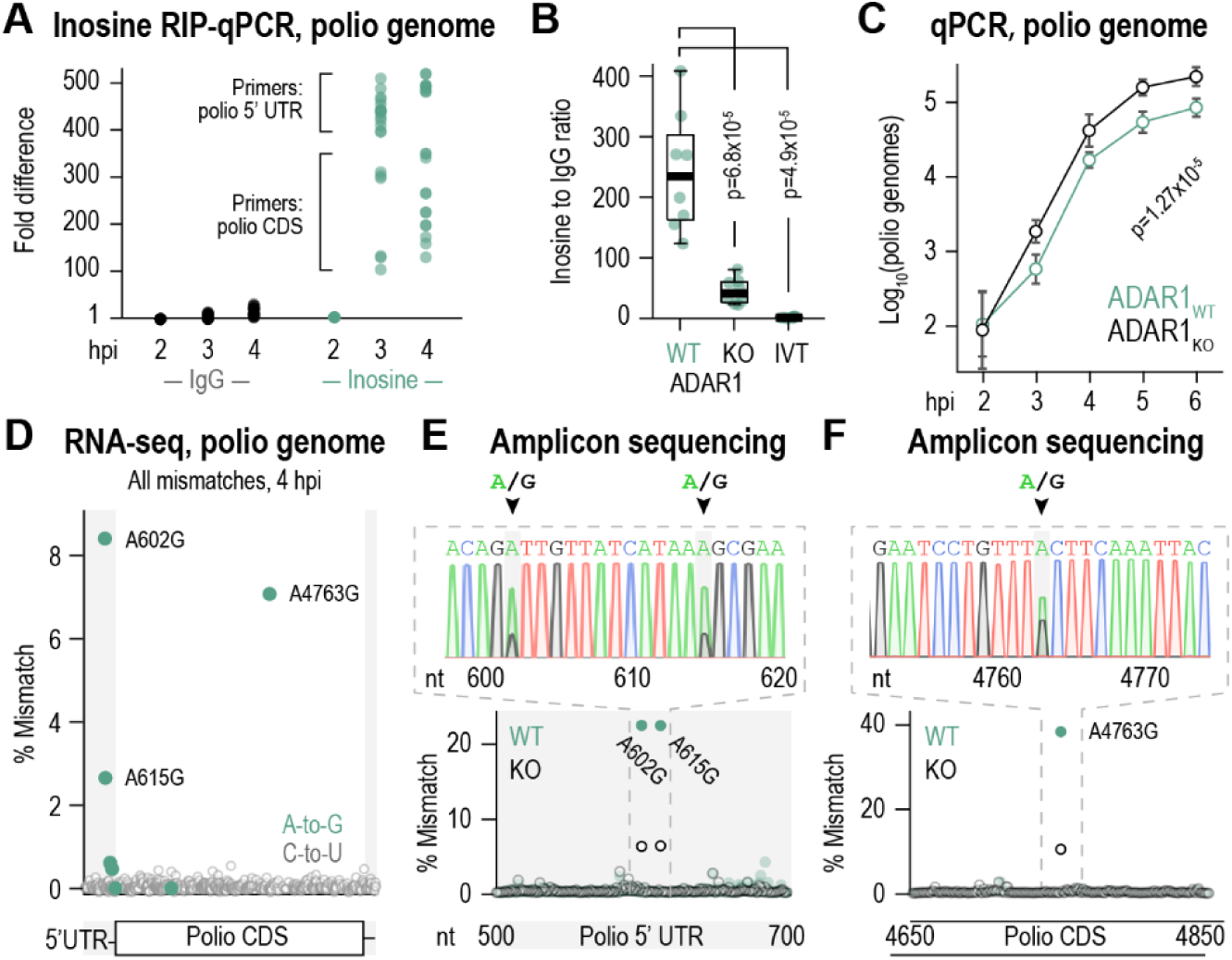
ADAR1 edits specific sites in polio vRNA. (**A**) Total RNA was isolated from infected cells (MOI=0.1) and incubated with anti-inosine beads to pull down inosine-containing RNA. Polio genomes were quantified by RT-qPCR (n=8 replicates, each amplified using 2 primer combinations). (**B**) Same as A, for RNA extracted from infected ADAR1_KO_ cells (MOI=0.1) and in vitro transcribed polio vRNA (IVT) at 4 hpi (n=4 replicates, each amplified using 2 primer combinations). Values are normalized to input RNA levels. p, Mann-Whitney p-value. (**C**) Total RNA was extracted from infected cells (MOI=0.1) at different timepoints. Polio genomes were quantified by RT-qPCR and normalized to GAPDH. Shown are means ±SD for each cell line (n=4). p, two-tailed Student T test p-value at 6 hpi. (**D**) All single nucleotide mismatches detected in polio vRNA by RNA-seq at 4 hpi (n=1), arranged by their position in the viral genome. Shaded areas represent UTRs. CDS, coding sequence. Shown is % mismatch of each site. (**E-F**) Total RNA was isolated from infected (MOI=0.1) ADAR1_WT_ or ADAR1_KO_ Huh7 cells and vRNA was amplified by PCR and analyzed by amplicon sequencing. The amplified regions span about 100 nt upstream and downstream of nucleotides 602/615 (E) and 4763 (F). Top, chromatograms showing mixed A/G nucleotides at 602, 615 and 4763 in ADAR1_WT_ cells. Bottom, % mismatch for every A-to-G substitution in the indicated region of the vRNA. Shown are representative chromatograms and mismatch plots (n=3, about 8,000 sequenced reads per sample).

Inosines form base pairing with cytosines and are therefore recognized as guanosines by RNA-dependent polymerases and other enzymes. Adenosine-to-guanosine (A-to-G) mismatches in RNA-seq data can be therefore be used as a proxy for ADAR editing^58^. To identify ADAR1 target sites in polio vRNA, we searched for RNA-seq reads that map to polio genome and harbor any mismatches from the reference sequence. At 4 hpi, we identified a total of 563 single nucleotide substitutions across the polio genome (Table S2). Most were cytosine-to-uracil (C-to-U) mismatches, consistent with nonspecific hyperediting by APOBEC cytosine deaminases^55^. These were distributed throughout the viral genome at low levels (Fig. 4D, gray, and Table S1). In contrast, A-to-G mismatches were limited to two sites in the 5’ UTR (genomic nucleotide positions 602, 615) and one site in the coding region (position 4763) but occurred at higher frequencies (Fig. 4D, green).

To confirm that these mismatches reflect sites of ADAR1 editing, we infected ADAR1_WT_ and ADAR1_KO_ cells (MOI=0.1), extracted RNA, and amplified vRNA using oligonucleotides that flank the three major A-to-G sites. Using amplicon sequencing, we detected the same A-to-G mismatches at positions 602, 615 and 4763 of the viral genome (Fig. 4E-F). These 3 positions were the most variable sites within a 200 nt region, and their mismatch rate was lower in ADAR1_KO_ cells (Fig. 4E-F). Residual A-to-G mismatches in ADAR_KO_ cells (Fig. 4E-F), similar to inosine content (Fig. 4B), are consistent with residual or redundant ADAR activity. Taken together, we conclude that ADAR1 edits polio vRNA on specific sites during infection.

### ADAR1 disrupts polio vRNA translation at both initiation and elongation

The adenosines at positions 602 and 615 of the vRNA map to the last dsRNA region of the IRES, known as domain VI, which harbors the ribosome landing site (Fig. 5A and S5A, adapted from^59^). Other mutations in domain VI reduce the efficiency of IRES-mediated translation in polio and related picornaviruses^19^. Therefore, we suspected that ADAR1 editing may reduce the efficiency of polio vRNA translation. To test this, we used a bicistronic reporter encoding two fluorescent proteins in tandem. A green fluorescent protein (EGFP) is produced by cap-dependent translation, whereas a red fluorescent protein (tdTomato) is translated from the polio IRES in the same mRNA^60^ (Fig. 5B). Red-to-green ratios measured by flow cytometry reflect the relative rate of IRES- vs. cap-mediated translation. A variant with guanosines at positions 602 and 615 (IRES^GG^) was included as an inosine mimetic that cannot be further edited by ADAR1. We transfected cells and measured red-to-green ratios for 10,000 cells in each sample. ADAR1 KO increased and overexpression (OE) decreased the efficiency of translation initiation from polio IRES^WT^ (Fig. 5C, left panel). Neither KO nor OE had significant effects on translation from IRES^GG^ (Fig. 5C, middle panel), which was less efficient than IRES^AA^ in ADAR1_WT_ cells (Fig. 5C, right panel). Therefore, ADAR1 inhibits internal translation initiation on polio vRNA, perhaps by affecting the structure of domain VI and reducing its ability to recruit initiating ribosomes.

**Fig. 5.**
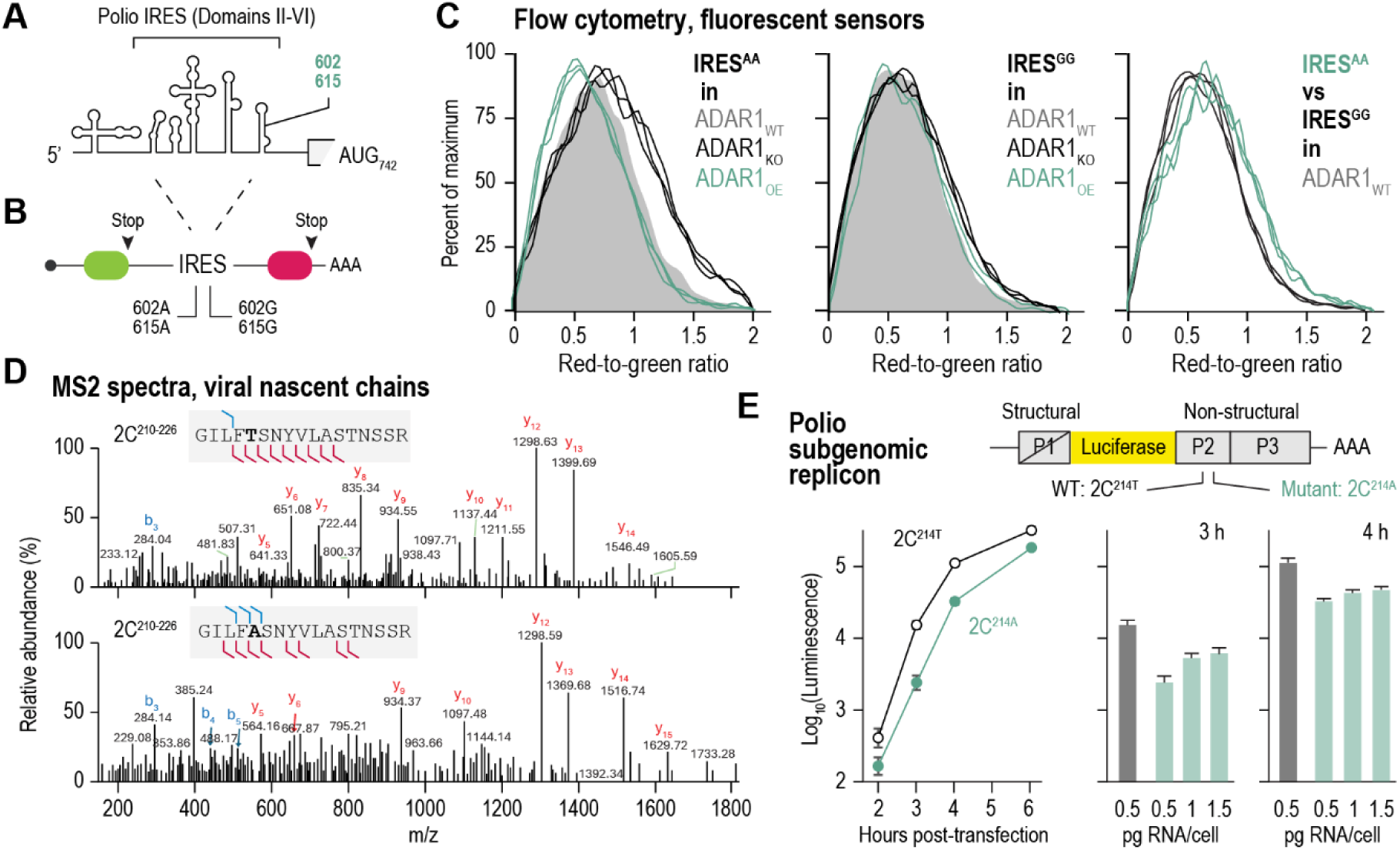
ADAR1 disrupts polio translation at both initiation and elongation. (**A**) Schematic secondary structure diagram of polio 5’ UTR, showing the positions of ADAR1 edits within Domain VI of the viral IRES. (**B**) Bicistronic translation sensor. The first open reading frame (ORF), encoding a green fluorescent protein (EGFP), is translated by cap-dependent initiation. The second ORF, encoding a red fluorescent protein (tdTomato), is translated by internal initiation from polio IRES. The IRES is either WT, with adenosines at positions 602 and 615 (IRES_AA_), or an A- to-G double mutant with guanosines at these positions (IRES_GG_). (**C**) Plasmids encoding the sensors from (B) were transfected into Huh7 cells expressing low, normal or high levels of ADAR1 (knockout, WT, overexpression; ADAR1_KO_, ADAR1_WT_, ADAR1_OE_) and red-to-green ratios were measured individually for each cell using flow cytometry at 24 hours post-transfection (∼10,000 cells per condition, n=3). Shown are population distributions of red-to-green ratios. Shifts to the right reflect higher and left lower relative efficiency of internal initiation from polio IRES. Left to right, IRES_AA_ transfection in cells expressing different levels of ADAR1; same, for IRES_GG_; and transfection of either IRES_AA_ or IRES_GG_ in ADAR1_WT_ cells. (**D**) To identify amino acid misincorporation in polio proteins, we isolated ribosome-associated nascent chains from infected cells using biotinylated puromycin and analyzed them by MS (n=3). Shown are MS2 m/z profiles for a viral peptide with two single amino acid variants: 2C^214T^ (WT, top) and 2C^214A^. (**E**) Polio luciferase replicons lacking structural proteins (P1) and harboring either adenosine or guanosine at genomic position 4763 were in vitro transcribed and transfected into Huh7 cells. Luciferase was measured at the indicated times post-transfection. Left, transfection of 0.5 pg replicon RNA per cell; right, transfection of 0.5 pg/cell WT versus 0.5-1.5 pg/cell 2C_214_ replicon (n=4).

Finally, we wondered how editing of polio vRNA at genomic position 4763 affects viral protein synthesis. Because ribosomes misread inosines as guanosines, ADAR1 editing of coding RNA can trigger miscoding^55,61^. We hypothesized that editing of position 4763 could make ribosomes miscode an ACT threonine codon as a GCT alanine codon, introducing a noncognate amino acid at position 214 of nonstructural protein 2C. To determine if this mutant viral protein is made in infected cells, we isolated polysomes at 4 hpi and used biotinylated puromycin to label and isolate nascent polypeptide chains. We then performed LC-MS/MS according to PUNCH-P, a protocol we developed specifically for nascent chain analysis^62^. As expected, polio was the most abundant nascent polypeptide at 4 hpi and peptides were distributed throughout the entire polyprotein (Fig. S5B-C and Table S3). Next, we performed a separate search allowing for single amino acid substitutions in viral nascent polypeptide chains. Remarkedly, only two sites of amino acid misincorporation were detected with high confidence and confirmed by MS2 (Fig. S5D and Table S3). One of those had either alanine or threonine at position 214 of polio 2C (Fig. 5D). This newly-synthesized error-containing peptide was not detected in uninfected cells or infected cells processed without biotinylated puromycin (Table S3), confirming that a mutant 2C^T214A^ protein is produced in infected cells.

Threonine at position 214 of polio 2C is conserved in other enteroviruses (Fig. S5E) but does not map close to the active site (Fig. S5F). To find if the 2C^T214A^ mutation affects viral fitness, we used a self-replicating polio-derived vRNA (replicon) model. This in vitro synthesized vRNA consists of the full viral genomic sequence, but regions coding for structural proteins were replaced with luciferase^63^. We mutated the replicon sequence to incorporate T214A in all 2C copies, synthesized vRNA in vitro, transfected Huh7 cells and monitored luminescence over time. The 2C^T214A^ mutant vRNA produced lower luminescence—a proxy for replication—at all timepoints tested from 2 to 6 h (Fig. 5E, left). Even when we transfected 2 or 3 times more of the 2C^T214A^ mutant vRNA, it still failed to reach the same luminescence as WT (Fig. 5E, right). We conclude that the T214A mutation affects polio 2C function and attenuates infection. Therefore, ADAR1 functions as a cellular barrier to polio infection by lowering the production of functional viral proteins through two distinct mechanisms.

## Discussion

Viruses are powerful tools for dissecting fundamental aspects of molecular cell biology, including RNA processing^2–4^. vRNA and encoded proteins interact with many of the host enzymes and molecular machines that participate in RNA processing. These host systems often play dual roles in infection; for example, translation is essential for producing viral proteins but also drives the production of innate antiviral factors^16^. Similarly, RNA editing can both enhance and suppress infection^55^. In +ssRNA viruses e.g. Zika, measles, and EV-D68, editing disrupts secondary RNA structures, helping vRNA evade detection by antiviral dsRNA sensors and thereby enhancing infection^64–66^. In contrast, editing of arenavirus vRNA reduces viral fitness by triggering a wide range of mutations in the encoded proteins^67^. Thus, RNA editing is a powerful evolutionary force in many viruses^68^.

Hyperediting by ADAR1 is common in many viruses, but reports on targeted or site-specific editing of vRNA remain limited^68^. In this study, we show that ADAR1 selectively and reproducibly edits three specific adenosines in the polio vRNA genome (Fig. 4D-F). Although details of the editing mechanisms remains to be elucidated, recent work has shown that ADAR1 activity is modulated by other RBPs^69^. Indeed, ILF2/3 and DHX9, which modulate ADAR1 activity^69^, were also recruited to polysomes during infection (Fig. 1E-F). It is possible that these and other RBPs may form complexes with ADAR1, guiding it to specific regions of the polio genome. Regardless of the mechanism that confers selective editing, our data shows that ADAR1 edits adenosines at positions 602 and 615 of polio vRNA, located within a stem-loop that harbors the ribosome landing pad of the IRES (Fig. S5A). ADAR1 also affects translation efficiency in other viruses, including Marburg, Ebola and EV-D68 viruses^64,70^. It is therefore tempting to speculate that ADAR1 reduces the efficiency of IRES-mediated translation, a hypothesis supported by our reporter assays (Fig. 5C).

Furthermore, our findings suggest that ADAR1 impairs polio protein synthesis through two distinct actions. It reduces the efficiency of IRES-mediated translation, lowering viral protein production; and introduces a site-specific miscoding event that results in mutant 2C proteins (Fig. 5D) with reduced replicative fitness (Fig. 5E). In hepatitis D virus (HDV), ADARs edit a UAG stop codon in the vRNA, enhancing its base pairing with tRNA-Trp(CCA) and triggering stop codon readthrough.

This produces a C-terminally extended viral protein that is essential for packaging^55^. Although ADAR editing is pro-viral in HDV, a similar mechanism may explain the T214A mutation in polio. Editing at position 4763 of the viral genome could weaken base pairing with the cognate tRNA-Thr(AGT) and enhance base pairing with tRNA-Ala(AGC), leading to a recurrent mutation in the polio 2C protein.

Picornavirus 2C proteins function in multiple stages of the viral life cycle, including replication and packaging^71^. 2C proteins are more conserved relative to other viral proteins and form AAA+ ATPase complexes^71^. Thr214 is located within a disordered region of 2C, adjacent to the intersubunit signaling (ISS) domain of the ATPase complex but distant from the ATPase active site (Fig. S5F). Mutations in polio 2C affect the susceptibility of the virus to certain replication inhibitors^72^. Structural modelling and docking studies suggest that Thr214 contributes to allosteric inhibitor binding at the ATPase domain but not the ATP binding site^73^. Further experiments are needed to determine how the T214A mutation affects 2C protein structure, post-translational modifications and oligomeric status. Given that single amino acid changes in polio can be both detrimental and beneficial, depending on environmental factors e.g. temperature^74^, the effects of the T214A mutation in 2C should also be evaluated under different physiological conditions. If this mutation confers a fitness advantage, ADAR1 editing of A4763 in polio vRNA could also be involved in viral adaptation.

Beyond their role in innate immunity, ADARs are important regulators of neuronal homeostasis^75^. It is therefore possible they are involved in the neurotropism and neurovirulence of polio. Previous studies have shown that A-to-G substitutions in the 5’ UTR of polio reduce the translation efficiency of viral genomes and limit neuroinvasion and neurovirulence^76,77^, although it is unclear whether these mutations are introduced by ADARs. A-to-G substitutions are also associated with reversion to neurovirulence of vaccine-derived strains^78^. Furthermore, neurons depend on ADAR-mediated miscoding: ADARs introduce a single coding error in GluA2, a glutamate receptor subunit, affecting its ion flow function^61^. Hypoediting of this site is linked to neurodegenerative diseases e,g, Amyotrophic Lateral Sclerosis (ALS) and Alzheimer’s^75,79^. In our experiments, “hypoediting” of polio vRNA in ADAR1_KO_ cells (Fig. 4E-F) increased vRNA production (Fig. 4C, S4). Future studies should assess how editing of polio vRNA by ADAR1 affects the neurovirulence of pathogenic and attenuated polio strains.

ADAR1 is part of a broader class of dsRNA sensors that recognize viral RNA duplexes and initiate innate immune responses^80^. Another member of this group, interferon-induced dsRNA-activated protein kinase (PKR or EIF2AK2), binds picornavirus IRES during translation^44^. Indeed, we observe that RNase I displaces PKR from polysomes translating polio vRNA (Table S1). This underscores a common vulnerability in +ssRNA viruses: while structured 5’ UTRs are essential for ribosome recruitment and translation, they also expose the vRNA to detection by dsRNA sensors. Therefore, innate immune activation by vRNA could be linked to its translation. The dynamic ensemble of RBPs that bind vRNA during translation may regulate infection by simultaneously controlling translation and innate immunity.

In conclusion, our findings suggest that ADAR1 functions as a polio restriction factor that critically limits the production of functional viral proteins. It does so by editing domain VI of the polio IRES, impairing translation initiation; and by editing the coding region of the polio genome, introducing a detrimental coding error. The experimental pipeline developed here could also be broadly applicable for identifying vRNA translation modulators and elucidating their roles in other viral pathogens.

## Resource availability

### Data and software availability

Sequencing data were deposited in SRA database under BioProject number PRJNA1257916. The mass spectrometry proteomics data have been deposited to the ProteomeXchange Consortium via the PRIDE^81^ partner repository with the dataset identifiers PXD064593 (RNase digestion) and PXD064639 (PUNCH-P).

## Acknowledgements

All infectious poliovirus work was done in Raul Andino’s biocontainment facility at UCSF. We thank Kuo-Feng Weng and Kathrin Leppek for critical reading of the manuscript and helpful discussions. This work was supported by funding from Chan Zuckerberg Biohub—SF and Chan Zuckerberg Initiative.

## Author contributions

D.K. performed experiments and analyzed data. K.L., F.M. acquired LC-MS/MS data. R.J.C. performed PUNCH-P MS data analysis to detect amino acid changes in polio nascent chains. J.E.E. supervised and provided funding for mass-spectrometry. J.P. performed polio 2C structural modeling. R.A. conceived the study, designed and performed experiments, analyzed data and wrote the manuscript.

## Supplementary Figure legends

**Figure S1.**
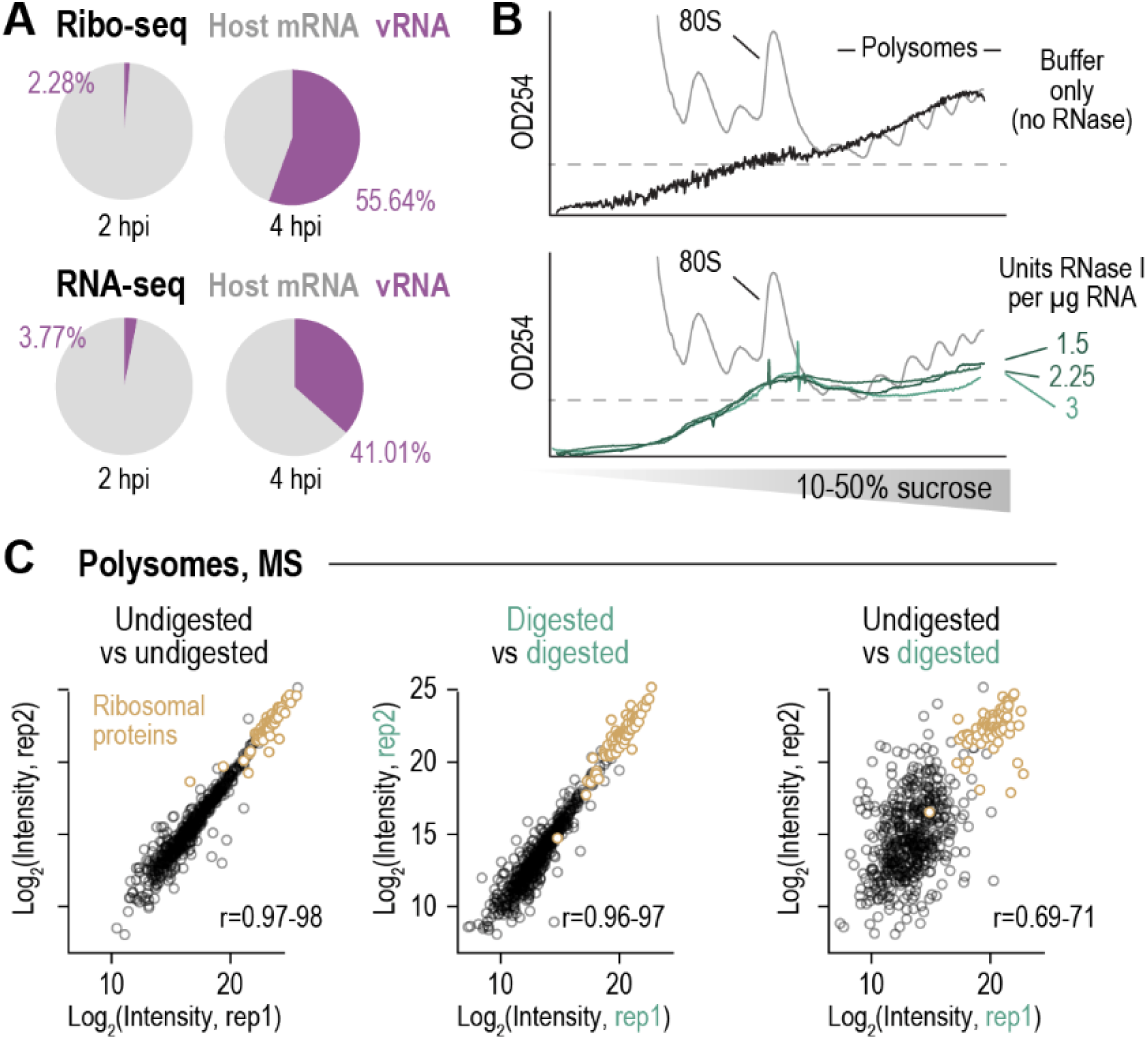
Ribosome profiling (ribo-seq), RNA sequencing (RNA-seq) and RNase digestion MS of polio infected cells, related to Figure 1. (**A**) Ribosome-protected (top) and total RNA (bottom) reads from ribo-seq (n=2) and RNA-seq (n=1) analysis of Huh7 cells infected with type I poliovirus (Mahoney strain) and collected at the indicated times. (**B**) Polysomes were extracted from uninfected Huh7 cells using 10-50% sucrose gradients and treated with RNase I at the indicated concentrations. Digested and control polysomes were pelleted through 60% sucrose cushions, and the pellets were resuspended and resolved again on 10-50% sucrose gradients. Shown are representative rRNA absorption plots for input cell lysate (gray), undigested polysome pellets (black), and RNase I-digested polysome pellets (green) (n=2). (**C**) Pairwise comparisons showing all individual proteins detected in two biological replicates from either digested or undigested polysomes, using iBAQ normalized intensities.

**Figure S2.**
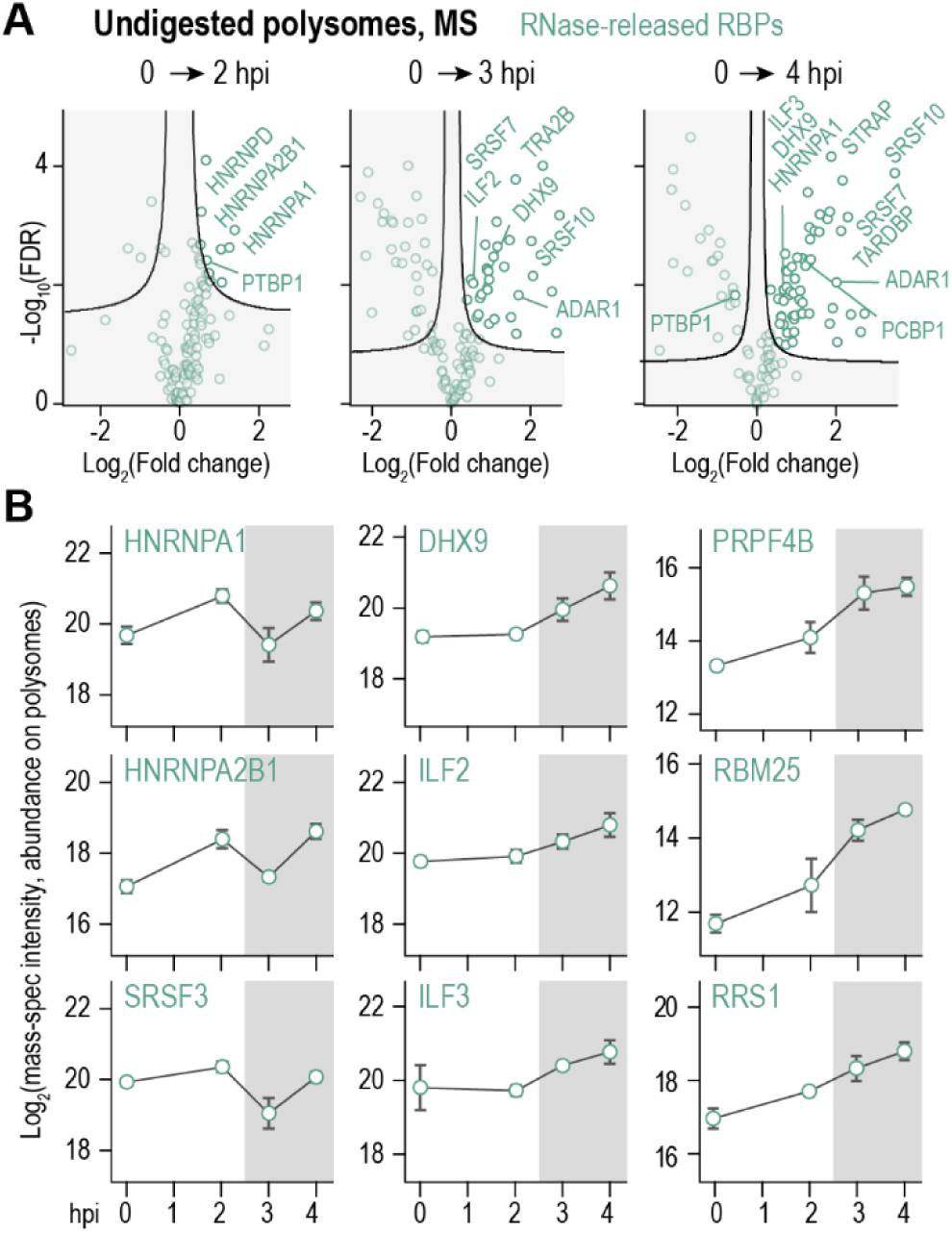
Temporal dynamics of polysomal RBPs during polio infection, related to Figure 2. (**A**) Pairwise comparison of RNase I-sensitive host RBPs detected on polysomes at either timepoint compared to uninfected controls (n=3 each). (**B**) Time course of polysome association for selected RBPs. Top to bottom, similar temporal patterns. Shown are means ±SD at each timepoint (n=3 each). Shading reflects global translation shutoff between 2 and 3 hpi.

**Figure S3.**
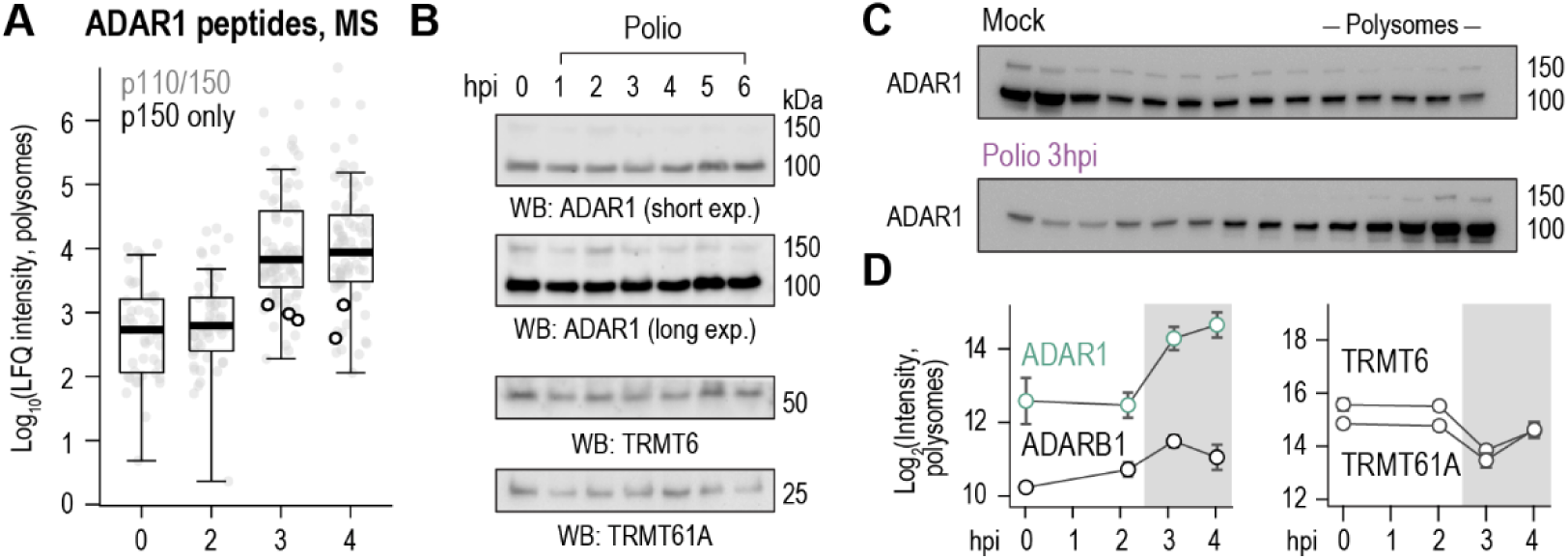
ADAR1 isoforms in polio infection, related to Figure 3. (**A**) peptide intensities normalized by label free quantification (LFQ) from 3 replicates of undigested polysomes. Peptides mapping only to the longer isoform of ADAR1, p150, are highlighted in circles. Peptides mapping to either p110 or p150 are in gray. (**B**) Huh7 cells were infected at MOI=5 and whole cell lysates were analyzed by immunoblotting at the indicated times. Blots are representative of n=2. (**C**) Longer exposure of gradient fractions from Fig. 3B, showing ADAR1 p150. (**D**) Polysome association timecourse for the indicated host proteins. Shown are mean ±SD (n=3).

**Figure S4.**
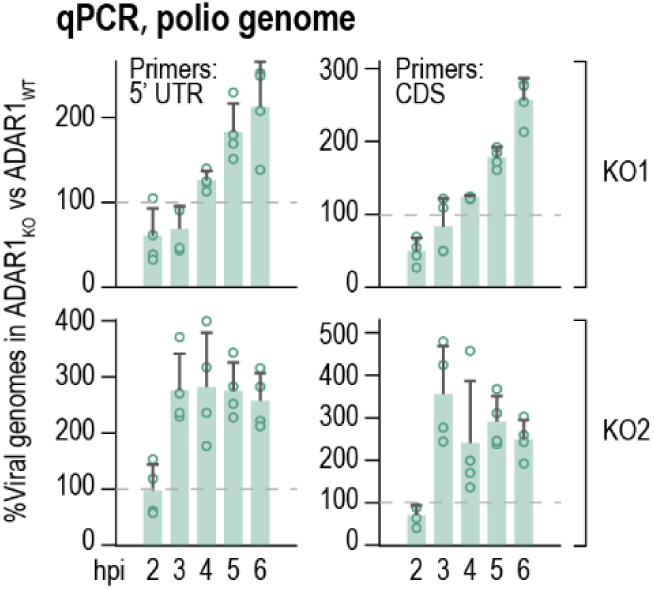
Effects of ADAR1 KO on polio genome editing and overall levels, related to Figure 4. ADAR1 KO Huh7 lines were generated using two targeting gRNAs (KO1 and KO2). Total RNA was extracted from infected cells (MOI=0.01) at different timepoints. Polio genomes were quantified by RT-qPCR using primers that amplify the 5’ UTR or CDS and normalized to GAPDH. Shown are means ±SD of percent change between ADAR1_KO_ and ADAR1_WT_ (n=4 each).

**Figure S5.**
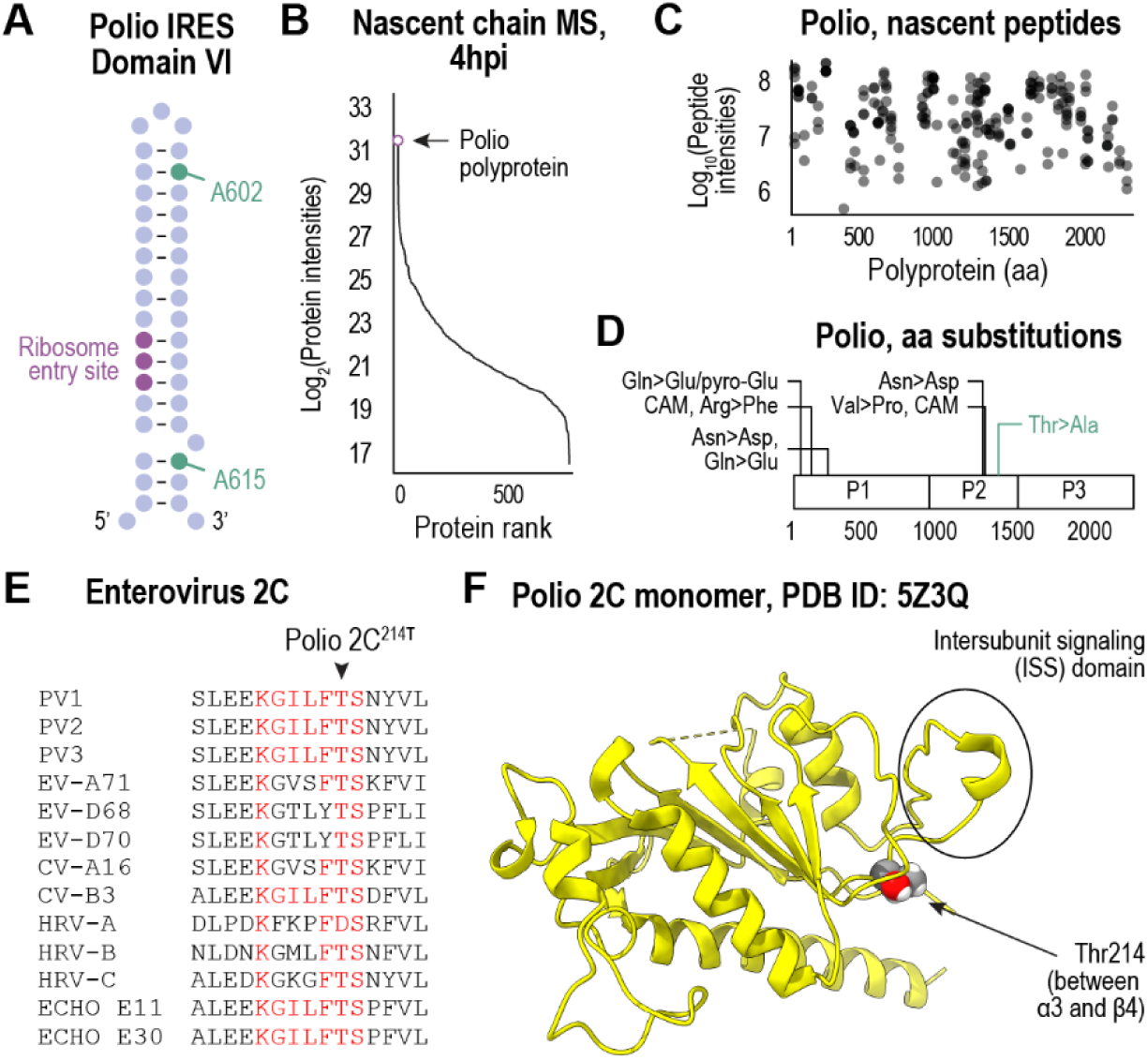
ADAR1 effects on translation initiation and elongation of polio, related to Figure 5. (**A**) Cartoon illustration of nucleotide base pairing in polio 5’ UTR domain VI. (**B-D**) PUNCH-P nascent chain MS analysis in infected (MOI=5) Huh7 cells. B, protein rank plot for all ribosome-associated nascent polypeptides detected. C, all ribosome-associated nascent polio peptides, plotted by their start position in the full-length viral polyprotein. (**D**) All amino acid (aa) substitutions detected in newly synthesized viral proteins. A threonine to alanine substitution at position 214 of polio 2C, corresponding to position 1341 of the full-length polyprotein, was detected and confirmed by MS2 (green). Other substitutions included asparagine deamidation to aspartic acid, a possible artifact of sample handling, and an arginine to phenylalanine substitution, which upon manual analysis could also be explained by trioxidation of a cysteine in the same peptide. Arg>Phe and carabamidomethyl (CAM) to trioxidation represent a similar mass shift of −9 Da, which is isobaric at the mass accuracy the data was acquired at. (**E**) Amino acid alignment for multiple enteroviruses showing conservation of Thr in position 214 of PV1 2C. (**F**) Ribbon diagram of Polio 2C AAA+ ATPase with position 214 indicated.

## Methods

### Key resource table

| REAGENT or RESOURCE | SOURCE | IDENTIFIER |
| --- | --- | --- |
| **Antibodies** |  |  |
| Mouse anti-puromycin | Millipore Sigma | Cat# MABE343 |
| Mouse anti-Polio 3D | Hybridoma<sup>82</sup> | N/A |
| Rabbit anti-RPLP1 | Abcam | Cat# ab121190 |
| Rabbit anti-ADAR1 | Cell Signaling | Cat# 14175S |
| Rabbit anti-TRMT6 | Proteintech | Cat# 16727-1-AP |
| Rabbit anti-TRMT61A | Proteintech | Cat# 31169-1-AP |
| Goat anti-Mouse IgG, HRP | Millipore Sigma | Cat# NA9310 |
| Goat anti-Rabbit IgG, HRP | Millipore Sigma | Cat# NA9340 |
| **Chemicals and kits** |  |  |
| Puromycin | Thermo Fisher | Cat# A1113803 |
| Lipofectamine 2000 | Thermo Fisher | Cat# 11668019 |
| Sequencing Grade Modified Trypsin | Promega | Cat# V5111 |
| Lys-C, Mass Spec Grade | Promega | Cat# VA1170 |
| ZipTip Pipette Tips | Millipore Sigma | Cat# ZTC18M960 |
| StrataClean Resin | Agilent | Cat# 400714 |
| RNase I | Thermo Fisher | Cat# AM2295 |
| SUPERase-In | Thermo Fisher | Cat# AM2694 |
| Complete EDTA-free protease inhibitor cocktail | Millipore Sigma | Cat# 11873580001 |
| Triton x-100 | Thermo Fisher | Cat # X100 |
| Sodium deoxycholate | Millipore Sigma | Cat # 30970 |
| Pellet paint co-precipitant | Millipore Sigma | Cat# 69049 |
| 4x Laemmli Sample Buffer | BioRad | Cat# 1610747 |
| Benzonase | Millipore Sigma | Cat# 70746 |
| TWEEN 20 | Millipore Sigma | Cat# 11332465001 |
| 4–20% Mini-PROTEAN TGX Precast Protein Gels | BioRad | Cat# 4561096 |
| Bovine Serum Albumin Fraction V | Millipore Sigma | Cat# 10735078001 |
| DMEM/F-12, GlutaMAX supplement | Thermo Fisher | Cat# 10565018 |
| Opti-MEM I Reduced Serum Medium, GlutaMAX Supplement | Thermo Fisher | Cat# 51985034 |
| DPBS, calcium, magnesium | Thermo Fisher | Cat# 14040133 |
| T4 RNA ligase II | New England Biolabs | Cat# M0242S |
| T4 polynucleotide kinase | New England Biolabs | Cat# M0201S |
| Universal miRNA Cloning Linker | New England Biolabs | Cat# S1315S |
| SuperScript III Reverse Transcriptase | Thermo Fisher | Cat# 18080093 |
| CircLigase ssDNA ligase | Epicentre | Cat# CL4111K |
| In Fusion Snap Assembly Master Mix | Takara | Cat # 638951 |
| Phusion High-Fidelity DNA Polymerase | New England Biolabs | Cat# M0530S |
| **Commercial assays** |  |  |
| Pierce BCA Protein Assay Kit | Thermo Fisher | Cat# 23225 |
| Pierce ECL Plus Western Blotting Substrate | Thermo Fisher | Cat# 32134 |
| MEGAscript T7 Transcription Kit | Thermo Fisher | Cat# AMB13345 |
| High Capacity cDNA Reverse Transcription Kit | Thermo Fisher | Cat# 4368814 |
| ONE-Glo EX Luciferase Assay System | Promega | Cat# E8110 |
| Itaq™ Universal SYBR | BioRad | Cat# 1725120 |
| **Experimental models: cell lines and viruses** |  |  |
| Huh7 human hepatoma | Raul Andino, UCSF | RRID: CVCL_0336 |
| Type I Polio (Mahoney) infectious clone | Raul Andino, UCSF<sup>83</sup> | prib(+)XpA |
| Type I Polio (Mahoney) luciferase replicon | Raul Andino, UCSF<sup>84</sup> | prib(+)Luc |
| **Software** |  |  |
| MaxQuant 1.6.7.0 |  | https://maxquant.org |
| Perseus 1.6.7.0 |  | https://maxquant.org/perseus |
| ProteinProspector 6.5.2 |  | https://prospector.ucsf.edu |
| ImageJ 1.52a |  | https://imagej.nih.gov/ij/ |

### Method details

#### Cell culture and virus propagation and infection

Huh7 human hepatocellular carcinoma cells were grown in DMEM/F-12 1:1 medium (Thermo Fisher) supplemented with 10% fetal bovine serum, 100 units/mL penicillin and 100 mg/mL streptomycin. Cells were grown at 37°C in a 5% CO_2_ incubator. To generate infectious poliovirus, we first synthesized viral RNA from an infectious clone. We linearized 10 µg of prib(+)XpA^83^ with 50 units of MluI for 2 h at 37°C. Linearized DNA was extracted with phenol:chloroform:isoamyl alcohol (25:24:1 v/v), and in vitro transcription was performed using MEGAscript T7 Transcription Kit (Thermo Fisher), according to the manufacturer instructions. 10 µg of transcribed viral RNA was electroporated into 4×10^6^ Huh7 cells in 0.4 mL PBS, 4 mm electroporation cuvette, at 270 V/960 µF (GenePulser, Bio-Rad). Cells were grown in T75 flasks for 48 h at 37°C, and subjected to three freeze-thaw cycles. Supernatants were centrifuged at 2500 x g for 5 min at 4°C to remove cell debris. Clarified supernatants containing passage 0 (p) virus were aliquoted and frozen at - 80°C. P0 viruses were amplified once in Huh7 cells to generate P1 virus stock, which was processed and frozen as above. P1 stock was used for all experiments. Titer was determined by plaque assays^26^. Infections were performed in Huh7 cells by adding virus stock to DMEM/F-12 without FBS and incubating cells with the inoculum for 45 min at 37°C, with occasional shaking. Inoculum was then replaced with DMEM/F-12/10% FBS for the remainder of the experiment.

#### Puromycylation, SDS-PAGE and immunoblotting

To label nascent chains in cultured cells, 1 µM puromycin (Thermo Fisher) was added to tissue culture media for 10 min at 37°C. For immunoblotting, adherent cells were washed twice with ice-cold PBS and lysed on plate with RIPA buffer (25 mM Tris-HCl pH=7.5, 150 mM NaCl, 1% NP-40, 0.5% Sodium deoxycholate) supplemented with 1 mM DTT, Complete EDTA-free protease inhibitor cocktail, and 50 units/mL benzonase (Millipore Sigma) to remove DNA. Lysis was performed on ice for 20 min and lysates were clarified by centrifugation for 10 min at 12,000 x g, 4°C. Protein concentration was determined by BCA assay (Thermo Fisher) and reconstituted in 1x Laemmli sample buffer (Bio-Rad) supplemented with fresh 1% 2-mercaptoethanol. 10-20 µg of each sample was resolved on 4-20% denaturing gels (Bio-Rad) and transferred to 0.2 µm PVDF membranes using a wet transfer apparatus. Membranes were blocked with 4% molecular biology grade BSA (Millipore Sigma) in tris-buffered saline supplemented with 0.1% Tween-20 (Millipore Sigma, TBST) for 1 h at RT then probed with primary antibodies for 2 h at RT. Primary antibodies were diluted 1:1000 in 4% BSA/TBST supplemented with 0.02% sodium azide. Secondary antibodies were diluted 1:10,000 in TBST. Western blot detection was done using ECL Plus Western Blotting Substrate (Thermo Fisher) and images were taken either by film radiography or BioRad GelDoc imager. Densitometry analysis was performed using ImageJ version 1.52a.

#### Ribo-seq and RNA-seq analyses

Ribo-seq and matching RNA-seq libraries were constructed essentially as described^85^. Briefly, infected cells were washed and harvested in ice-cold PBS (with calcium and magnesium, Thermo Fisher) by scraping and pelleted by centrifugation at 1000 x g for 5 min at 4°C. Pellets were resuspended in polysome buffer (25 mM Tris-HCl pH=7.5, 150 mM NaCl, 10 mM MgCl_2_, 2 mM dithiothreitol (DTT) and Complete EDTA-free protease inhibitor cocktail (Millipore Sigma)). Triton X-100 and sodium deoxycholate (Milipore Sigma) were added to a final concentration of 1%. After 20 min on ice, the samples were centrifuged at 20,000 x g for 10 min at 4°C to remove cell debris. Lysates were clarified by centrifugation at 20,000 x g for 10 min at 4°C and fractionated on 10-50% sucrose gradients (see below). Polysome fractions were pooled and diluted 1:1 v/v in polysome buffer (25 mM Tris-HCl pH=7.5, 150 mM NaCl, 10 mM MgCl_2_, 2 mM dithiothreitol (DTT) and Complete EDTA-free protease inhibitor cocktail (Millipore Sigma)). RNA concentration was measured by nanodrop and 1.5 units Ambion RNase I (Thermo Fisher) were added per 1 µg total RNA. Digests were incubated for 1 h at 25°C with shaking at 1000 RPM, and digestion was terminated by addition of 200 U Superase-In (Thermo Fisher). RNA was extracted using phenol:chloroform:isoamyl alcohol and size selected for fragments 16-34 nt using an 15% urea-PAGE. Chemically fragmented input RNA was size selected as well. rRNA depletion was performed using Ribo-Zero rRNA Removal Kit (Illumina), according to the manufacturer’s instructions. All downstream steps were performed as described^85^. Libraries were sequenced on a HiSeq 4000 (Illumina). After demultiplexing, sequencing reads were trimmed of adaptor sequences and quality filtered using cutadapt (-a CTGTAGGCACCATCAAT -m1 -q20). Ribosomal RNA was removed using Bowtie2 (--un). Remaining reads were aligned to poliovirus genome (GenBank V01149.1 with a single base substitution U2133C found in prib(+)XpA) using Hisat2 (--trim5 1). Read count matrices were generated using featureCounts and BigWig files were generated using bamCoverage and visualized on Integrated Genome Viewer. Mismatch analysis was performed using Call variants with LoFreq (Galaxy Version 2.1.5+galaxy3)^86^.

#### Polysome profiles using sucrose gradients

1-5×10^7^ Huh7 cells were harvested by scraping in ice-cold PBS (with calcium and magnesium, Thermo Fisher), centrifuged 1000 x g for 5 min at 4°C, resuspended in PBS and centrifuged again. Cell pellets were flash frozen in liquid nitrogen. On the day of experiment, pellets were thawed on ice and resuspended in 200 µl polysome buffer (25 mM Tris-HCl pH=7.5, 150 mM NaCl, 10 mM MgCl_2_, 2 mM dithiothreitol (DTT) and Complete EDTA-free protease inhibitor cocktail (Millipore Sigma)). Triton X-100 and sodium deoxycholate (Milipore Sigma) were added to a final concentration of 1%. After 20 min on ice, the samples were centrifuged at 20,000 x g for 10 min at 4°C to remove cell debris. Clarified lysates were loaded on 10-50% sucrose gradients in polysome buffer and subjected to ultracentrifugation at 36,000 rpm in an SW41.Ti swinging bucket rotor (Beckman Coulter) for 150 min at 4°C. Equal volume fractions were collected using Gradient Station (BioComp) with continuous monitoring of rRNA at UV254. For immunoblot analyses, each fraction was incubated with 7 µl StrataClean beads (Agilent) for 16 h at 4°C with constant tumbling. Beads were pelleted by centrifugation at 1000 x g for 5 min at 4°C and the supernatant was removed. Proteins were eluted from beads in 30 µl 2x Laemmli buffer, and incubated at 95°C for 5 min.

#### Polysome isolation with RNase treatment

Polysome fractions from gradients were diluted 1:1 with polysome buffer and RNA concentration was determined by nanodrop. 1.5 units Ambion RNase I (Thermo Fisher) were added per 1 µg total RNA and digestion was performed for 1 h at 25°C with shaking at 1000 RPM, then terminated by addition of 200 U Superase-In (Thermo Fisher). Digested and undigested polysomes were loaded on 1 mL of 66% sucrose and subjected to ultracentrifugation at 50,000 rpm in a 70.1 Ti fixed angle rotor (Beckman Coulter) for 16 h at 4 °C. Pellets were washed once by gently dispensing and removing 1 mL of ice-cold polysome buffer before finally resuspending in 100 µl polysome buffer. Proteins were extracted using methanol-chloroform precipitation. 400 µl methanol, 100 µl chloroform and 350 µl water were added sequentially to each 100 µl sample, followed by centrifugation at 14,000 x g for 5 min at room temperature. The top phase was removed and the protein interphase was precipitated by addition of 400 µl methanol, followed by centrifugation at 14,000 g for 5 min at room temperature. Pellets were air dried and resuspended in 8M urea, 25 mM ammonium bicarbonate (pH 7.5).

#### Puromycin-associated nascent chain proteomics (PUNCH-P)

∼2×10^7^ Huh7 cells were infected at MOI=5, in triplicates, and harvested at 4 hpi as above. Polysome isolation from infected and uninfected cells, nascent chain labeling using biotinylated puromycin and streptavidin affinity capture were according to PUNCH-P^62^. To control for nonspecific binding to streptavidin beads, a similar number of infected cells were processed as above without biotinylated puromycin.

#### LC-MS/MS sample preparation and data acquisition

For polysome analyses, protein concentration was determined by BCA (Thermo Fisher) and 1-2 µg total protein were subjected to reduction and alkylation by incubation with 10 mM DTT for 1 h at room temperature followed by 5 mM iodoacetamide for 1h at room temperature, in the dark. For PUNCH-P, nascent chains bound to streptavidin beads were incubated with 10 mM DTT for 1h at room temperature followed by 5 mM iodoacetamide for 1h at room temperature, in the dark. All samples were adjusted to 2M urea in 25 mM ammonium bicarbonate (pH 7.5) and sequencing-grade trypsin (Promega) was added at 1:50 enzyme to protein ratio (polysomes) or 1 µg (PUNCH-P). Digests were incubated overnight at 25°C and acidified with 0.1% trifluoroacetic acid. For polysome samples, peptides were desalted with μC18 Ziptips (Millipore Sigma), dried and resuspended in 10 μL 0.1% formic acid in water. For PUNCH-P samples, peptides were desalted with in-house styrenedivinylbenzene reversed phase sulfonate packed stagetips^87^, dried and resuspended in 2% acetonitrile/0.1% TFA.

Data acquisition, polysomes: peptides were analyzed on a Velos Pro Elite Orbitrap mass spectrometer (Thermo Fisher) coupled online to a nanoAcquity UPLC system equipped with an EASY-Spray nanoESI ion source (Thermo Fisher). Peptides were separated by capillary reverse phase chromatography on an EASY-Spray column (75 μm x 15 cm column packed with 3μm, 100 Å PepMap C18 resin) at 2% B (0.1% formic acid in acetonitrile) for 20 min at a flow rate of 600nl/min. Peptides were separated at 400 nL/min using a gradient from 2% to 25% B over 220 min followed by a second gradient from 25% to 37% B over 8 minutes and then a column wash at 75% B and reequilibration at 2% B. Precursor scans were acquired in the Orbitrap analyzer (300-1800 m/z, resolution: 60,000@400 m/z, AGC target: 2e6). The top 6 most intense, doubly charged or higher ions were isolated (4 m/z isolation window), subjected to high-energy collisional dissociation (27.5 NCE), and the product ions measured in the orbitrap analyzer (15,000@400 m/z, AGC target: 9e4).

Data acquisition, PUNCH-P: peptides were analyzed on a Fusion Lumos mass spectrometer (Thermo Fisher) equipped with a Thermo EASY-nLC 1200 LC system (Thermo Fisher). Peptides were separated by capillary reverse phase chromatography on a 25 cm column (75 µm inner diameter, packed with 1.7 µm C18 resin, AUR3-25075C18, Ionopticks, Victoria Australia). Peptides were introduced into a Fusion Lumos using a two-step linear gradient with 3–27 % buffer B (0.1% (v/v) formic acid in 80% (v/v) acetonitrile) for 52.5 min followed by 27-40 % buffer B for 14.5 min at a flow rate of 300 nL/min. Column temperature was maintained at 50°C throughout the procedure. Data was acquired in top speed data dependent mode with a duty cycle time of 1 s. Full MS scans were acquired in the Orbitrap mass analyzer (FTMS) with a resolution of 120,000 (FWHM) and m/z scan range of 375-1500 m/z. Selected precursor ions were subjected to fragmentation using higher-energy collisional dissociation (HCD) with quadrupole isolation window of 0.7 m/z, and normalized collision energy of 31%. HCD fragments were analyzed in the Ion Trap mass analyzer (ITMS) set to Turbo scan rate. Fragmented ions were dynamically excluded from further selection for a period of 60 seconds. The AGC target was set to 1,000,000 and 10,000 for full FTMS and ITMS scans, respectively. The maximum injection time was set to Auto for both full FTMS scans and ITMS scans.

#### MS data processing

Raw MS data for polysomes were processed using MaxQuant version 1.6.7.0^88^. MS/MS spectra searches were performed using the Andromeda search engine^89^ against the forward and reverse human proteome (UP000005640) and type I poliovirus (P03300, both downloaded November 25, 2020). Cysteine carbamidomethylation was chosen as fixed modification and methionine oxidation and N-terminal acetylation as variable modifications. Precursor and fragment mass tolerances were set at 4.5 and 20 ppm, respectively. Maximum allowed false discovery rate (FDR) was <0.01 at both the peptide and protein levels, based on a standard target-decoy database approach. The “calculate peak properties” and “match between runs” options were enabled.

Raw MS data for PUNCH-P were processed using Protein Prospector version 6.5.2. MS/MS spectra were searched against forward and sequence-randomized human proteome (UP000005640) and type I poliovirus (P03300). Cysteine carbamidomethylation was chosen as a fixed modification and methionine oxidation, protein N-terminal acetylation, and pyroglutamate formation from peptide N-terminal glutamine as variable modifications. The search was performed under ‘Homology’ setting (under Matrix Modifications), which allows for substitution of any single amino acid for another. Precursor and fragment tolerances were set at 5ppm and 0.4 Da, respectively. FDR was <0.01 at the peptide level and all modified peptide results were manually assessed.

#### Statistical testing

Statistical tests were performed with Perseus version 1.6.7.0 using either ProteinGroups or Peptides output tables from MaxQuant. Proteins identified in the reverse dataset and proteins only identified by site were filtered out. Intensity-based absolute quantification (iBAQ) was used to estimate absolute protein abundance. Two-sided Student’s t-test with a permutation-based FDR of 0.01 and S0^90^ of 0.1 with 250 randomizations was used to determine statistically significant differences between grouped replicates. Categorical annotations were based on Gene Ontology Biological Process (GOBP), Molecular Function (GOMF) and Cellular Component (GOCC), as well as protein complex assembly by CORUM.

#### Quantitative Real-Time PCR (qRT-PCR)

Total RNA was extracted using Trizol (Thermo Fisher) according to the manufacturer instructions. cDNA was synthesized from 100 ng RNA sample using High Capacity cDNA Reverse Transcription Kit (Thermo Fisher). qRT-PCR analysis was performed using Itaq Universal SYBR (Biorad). Primers used: GAPDH (Fwd 5’ AGGTCGGAGTCAACGGAT 3’, Rev 5’ TCCTGGAAGATGGTGATG 3’), Polio genome 5’ UTR, nt 503-702 (Fwd 5’ TTGGCCTGTCGTAACGCG 3’, Rev 5’ ACTTAGAGTAAACACACTCAATGGAGCG 3’), Polio genome CDS, nt 4663-4862 (Fwd 5’ AAACCCAGATGGTGCGGAC 3’, Rev 5’ CCATGTCGAACGCAAAGCG 3’).

#### Inosine immunoprecipitation

Huh7 cells were infected at MOI=0.1, washed with ice-cold PBS, and harvested in Trizol. Total RNA was extracted according to the manufacturer instructions, resuspended in water, and 10 µg was denatured at 80°C for 2 min followed by cooling on ice. To prepare antibody-coated beads, 5 µl protein A/G magnetic beads (per sample) were washed twice with 125 µl IP buffer (10 mM Tris pH 7.5, 0.1% IGEPAL CA-630, 150 mM NaCl). 125 ng of either anti-inosine or IgG control were added in 125 µl IP buffer and tumbled overnight at 4°C with shaking at 1000 RPM on a thermomixer. For each sample, 5 µg denatured RNA + 0.5 µl RNaseIn were added to either inosine or IgG coated beads and incubated at 4°C for 3 h with shaking as above. Beads were washed twice by incubating with 250 µl IP buffer for 10 min at 4°C, with shaking as above. Next, two more wash steps were completed as above using 250 µl low salt buffer (same as IP buffer only with 50 mM NaCl). Two more washes were performed as above using 250 µl high salt buffer (same as IP buffer only with 500 mM NaCl). Beads were resuspended in 50 µl RLT buffer (Qiagen) and incubated for 5 min at RT, with shaking as above. Eluates were transferred to new tubes and precipitated by 1.5 µl glycoblue and 2x v/v isopropanol, followed by overnight incubation at −20°C. Precipitation reactions were centrifuged at maximum speed for 30 min at 4°C, and pellets were washed with ice-cold 75% ethanol, followed by spindown for 5 min at 7500 g. Pellets were air dried and resuspended in 15 µl water. cDNA synthesis and RT-qPCR were performed as described above.

#### Targeted amplicon sequencing

Regions of Polio genome were amplified using primers designed to amplify about 100 nt region up and downstream of each potential ADAR editing site (see PCR primers), using a Q5 2X Hot Start HiFi master mix. 20 µl of each PCR reaction were transferred to fresh tubes and 36 µl AMPure XP Beads were added to each (representing a 1.8x v/v ratio). After vigorous pipetting, samples were incubated for 5 min at RT. On a magnetic separator, beads were washed three times with 200 µl 70% ethanol. Beads were resuspended in 40 µl water. After vigorous pipetting, samples were incubated for 2 min at RT. Beads were returned to the magnetic rack and supernatants were transferred to fresh tubes. cDNA was quantified by nanodrop and submitted for Premium PCR sequencing by Plasmidsaurus. About 8000 reads were analyzed for each sample.

#### Tandem fluorescence sensors for Polio IRES

A tandem fluorescent reporter of Polio IRES activity was reported elsewhere^60^. In this bicistronic plasmid, eGFP is translated by cap-dependent initiation and tdTomato is translated by cap-independent initiation from Polio IRES. A-to-G substitutions corresponding to positions 602 and 615 were generated by In-Fusion cloning (Takara) using Fwd primer 5’TCATAAGGCGAATTGGATAGGATC3’ and Rev primer 5’TAACAACCTGTGATTGTCACCATAAG3’. For transfections, Huh7 cells were plated in 12 wells at 1×10^5^ cells/well. The following day, 1.25 µg reporter plasmid were combined with 2.5 µl Lipofectamine 2000 in a total of 100 µl OptiMEM (Thermo Fisher), according to the manufacturer instructions. For co-transfection with an ADAR expression plasmid, 800 ng reporter plasmid plus 400 ng ADAR plasmid were combined with 2.5 ul Lipofectamine 2000 in a total of 100 ul OptiMEM. 100 µl transfection mixtures were diluted in 1.5 mL DMEM/10% FBS and added to cells. Media was replaced 6 h later with 1.5 mL DMEM/10% FBS. At 24 h post-transfection, cells were washed with PBS, dissociated with trypsin and diluted in 1 mL PBS. 10,000 were counted for every sample using a Cytoflex S (Beckman Coulter). Per cell red-to-green fluorescence ratios were calculated from the singlet population, gated to exclude cells without fluorescence.

#### Polio luciferase replicon

To form a luciferase replicon, an infectious clone of Polio was modified to replace capsid with firefly luciferase^84^. A-to-G substitution at the position corresponding to 4763 was generated by in-fusion cloning using Fwd primer 5’AATCCTGTTTGCTTCAAATTACG3’ and Rev primer 5’CCTTTCTCCTCCAGG3’. RNA coding for Polio-luciferase was generated as described above for generating the infectious virus. Plasmid DNA was digested by MluI for 2 h at 37°C, and linearized DNA was extracted using phenol:chloroform:isoamyl alcohol (25:24:1 v/v), followed by in vitro transcription using MEGAscript T7 Transcription Kit (Thermo Fisher), according to the manufacturer instructions. Huh7 cells were plated in black-wall 96 wells at 2.5×10^4^ cells/well. The following day, 10 ng RNA was mixed with 0.1 µl Lipofectamine 2000 in a total of 2.5 µl OptiMEM, according to manufacturer instructions. At the indicated times post-transfection, luminescence was measured by ONE-Glo EX Luciferase Assay System (Promega) using SpectraMax M3 (Molecular Devices) with 0.5 sec integration time.

#### Polio 2C structural model

A single protomer of the PV 2C ATPase crystal structure (Chain A, PDB ID: 5Z3Q) was displayed in ribbons, with T214 displayed in spheres, using ChimeraX 1.8^91^. To estimate the position of the intersubunit signaling (ISS) domain in hexameric PV 2C ATPase, a model was generated by superimposing the Chain A protomer of PV 2C ATPase (PDB ID:5Z3Q) onto each protomer of hexameric mitochondrial inner membrane AAA+ protease YME1 (PDB ID:6AZ0), using Matchmaker in ChimeraX 1.8 with default settings (Needleman-Wunsch alignment, BLOSUM-62 matrix, 0.3 secondary structure weighting).

## References

1. Iselin, L., Palmalux, N., Kamel, W., Simmonds, P., Mohammed, S., and Castello, A. (2022). Uncovering viral RNA–host cell interactions on a proteome-wide scale. Trends Biochem. Sci. 47, 23–38. 10.1016/j.tibs.2021.08.002.

2. Bermudez, Y., Hatfield, D., and Muller, M. (2024). A Balancing Act: The Viral–Host Battle over RNA Binding Proteins. Viruses 16, 474. 10.3390/v16030474.

3. Hanson, W.A., Romero Agosto, G.A., and Rouskin, S. (2024). Viral RNA Interactome: The Ultimate Researcher’s Guide to RNA–Protein Interactions. Viruses 16, 1702. 10.3390/v16111702.

4. Lal, A., Galvao Ferrarini, M., and Gruber, A.J. (2022). Investigating the Human Host-ssRNA Virus Interaction Landscape Using the SMEAGOL Toolbox. Viruses 14. 10.3390/v14071436.

5. Polio | Polio | CDC https://www.cdc.gov/polio/index.html.

6. O’Grady, M., and Bruner, P.J. (2024). Polio vaccine. In StatPearls [Internet] (StatPearls Publishing).

7. Sabin, A.B. (1985). Oral Poliovirus Vaccine: History of Its Development and Use and Current Challenge to Eliminate Poliomyelitis from the World. J. Infect. Dis. 151, 420–436. 10.1093/infdis/151.3.420.

8. Gromeier, M., Alexander, L., and Wimmer, E. (1996). Internal ribosomal entry site substitution eliminates neurovirulence in intergeneric poliovirus recombinants. Proc. Natl. Acad. Sci. 93, 2370–2375. 10.1073/pnas.93.6.2370.

9. Gutiérrez, A.L., Denova-Ocampo, M., Racaniello, V.R., and del Angel, R.M. (1997). Attenuating mutations in the poliovirus 5’ untranslated region alter its interaction with polypyrimidine tract-binding protein. J. Virol. 71, 3826–3833. 10.1128/jvi.71.5.3826-3833.1997.

10. Pilipenko, E. V, Viktorova, E.G., Guest, S.T., Agol, V.I., and Roos, R.P. (2001). Cell-specific proteins regulate viral RNA translation and virus-induced disease. EMBO J. 20, 6899–6908. 10.1093/emboj/20.23.6899.

11. Ochs, K., Zeller, A., Saleh, L., Bassili, G., Song, Y., Sonntag, A., and Niepmann, M. (2003). Impaired Binding of Standard Initiation Factors Mediates Poliovirus Translation Attenuation. J. Virol. 77, 115–122. 10.1128/JVI.77.1.115-122.2003.

12. Guest, S., Pilipenko, E., Sharma, K., Chumakov, K., and Roos, R.P. (2004). Molecular Mechanisms of Attenuation of the Sabin Strain of Poliovirus Type 3. J. Virol. 78, 11097–11107. 10.1128/JVI.78.20.11097-11107.2004.

13. Avanzino, B.C., Jue, H., Miller, C.M., Cheung, E., Fuchs, G., and Fraser, C.S. (2018). Molecular mechanism of poliovirus Sabin vaccine strain attenuation. J. Biol. Chem. 293, 15471–15482. 10.1074/jbc.RA118.004913.

14. Bedard, K.M., and Semler, B.L. (2004). Regulation of picornavirus gene expression. Microbes Infect. 6, 702–713. 10.1016/j.micinf.2004.03.001.

15. Hinnebusch, A.G., and Lorsch, J.R. (2012). The Mechanism of Eukaryotic Translation Initiation: New Insights and Challenges. Cold Spring Harb. Perspect. Biol. 4, a011544–a011544. 10.1101/cshperspect.a011544.

16. Rozman, B., Fisher, T., and Stern-Ginossar, N. (2022). Translation—A tug of war during viral infection. Mol. Cell. 10.1016/j.molcel.2022.10.012.

17. Mizuguchi, H., Xu, Z., Ishii-Watabe, A., Uchida, E., and Hayakawa, T. (2000). IRES-dependent second gene expression is significantly lower than cap-dependent first gene expression in a bicistronic vector. Mol. Ther. 1, 376–382. 10.1006/mthe.2000.0050.

18. Andreev, D.E., Hirnet, J., Terenin, I.M., Dmitriev, S.E., Niepmann, M., and Shatsky, I.N. (2012). Glycyl-tRNA synthetase specifically binds to the poliovirus IRES to activate translation initiation. Nucleic Acids Res. 40, 5602–5614. 10.1093/nar/gks182.

19. Sweeney, T.R., Abaeva, I.S., Pestova, T. V, and Hellen, C.U.T. (2014). The mechanism of translation initiation on Type 1 picornavirus IRESs. EMBO J. 33, 76–92. 10.1002/embj.201386124.

20. Svitkin, Y. V., Maslova, S. V., and Agol, V.I. (1985). The Genomes of attenuated and virulent poliovirus strains differ in their in vitro translation efficiencies. Virology 147, 243–252. 10.1016/0042-6822(85)90127-8.

21. Pilipenko, E. V, Pestova, T. V, Kolupaeva, V.G., Khitrina, E. V, Poperechnaya, A.N., Agol, V.I., and Hellen, C.U. (2000). A cell cycle-dependent protein serves as a template-specific translation initiation factor. Genes Dev. 14, 2028–2045.

22. Kauder, S.E., and Racaniello, V.R. (2004). Poliovirus tropism and attenuation are determined after internal ribosome entry. J. Clin. Invest. 113, 1743–1753. 10.1172/JCI21323.

23. Semler, B.L. (2004). Poliovirus proves IRES-istible in vivo. J. Clin. Invest. 113, 1678–1681. 10.1172/JCI22139.

24. Fernández-García, L., and Garcia-Blanco, M.A. (2025). Host RNA-binding proteins and specialized viral RNA translation mechanisms: Potential antiviral targets. Antiviral Res. 237, 106142. 10.1016/j.antiviral.2025.106142.

25. Yeh, M. Te, Bujaki, E., Smith, M., Weiner, A.J., Bandyopadhyay, A., Van Damme, P., De Coster, I., Revets, H., Macadam, A., and Andino, R. (2019). A New Live-Attenuated Poliovirus Vaccine Constructed by Rational Design Improves Safety by Preventing Reversion to Virulence. SSRN Electron. J. 10.2139/ssrn.3400856.

26. Aviner, R., Li, K.H., Frydman, J., and Andino, R. (2021). Cotranslational prolyl hydroxylation is essential for flavivirus biogenesis. Nature 596, 558–564. 10.1038/s41586-021-03851-2.

27. Gradi, A., Svitkin, Y. V., Imataka, H., and Sonenberg, N. (1998). Proteolysis of human eukaryotic translation initiation factor eIF4GII, but not eIF4GI, coincides with the shutoff of host protein synthesis after poliovirus infection. Proc. Natl. Acad. Sci. 95, 11089–11094. 10.1073/pnas.95.19.11089.

28. Gerashchenko, M. V., and Gladyshev, V.N. (2017). Ribonuclease selection for ribosome profiling. Nucleic Acids Res. 45, e6–e6. 10.1093/nar/gkw822.

29. Bertolini, M., Fenzl, K., Kats, I., Wruck, F., Tippmann, F., Schmitt, J., Auburger, J.J., Tans, S., Bukau, B., and Kramer, G. (2021). Interactions between nascent proteins translated by adjacent ribosomes drive homomer assembly. Science (80-.). 371, 57–64. 10.1126/science.abc7151.

30. Fusco, C.M., Desch, K., Dörrbaum, A.R., Wang, M., Staab, A., Chan, I.C.W., Vail, E., Villeri, V., Langer, J.D., and Schuman, E.M. (2021). Neuronal ribosomes exhibit dynamic and context-dependent exchange of ribosomal proteins. Nat. Commun. 12, 6127. 10.1038/s41467-021-26365-x.

31. Norris, K., Hopes, T., and Aspden, J.L. (2021). Ribosome heterogeneity and specialization in development. WIREs RNA 12. 10.1002/wrna.1644.

32. Pöyry, T., Stoneley, M., and Willis, A.E. (2020). Should I Stay or Should I Go: eIF3 Remains Ribosome Associated and Is Required for Elongation. Mol. Cell 79, 539–541. 10.1016/j.molcel.2020.07.025.

33. Stein, K.C., Kriel, A., and Frydman, J. (2019). Nascent Polypeptide Domain Topology and Elongation Rate Direct the Cotranslational Hierarchy of Hsp70 and TRiC/CCT. Mol. Cell 75, 1117–1130.e5. 10.1016/j.molcel.2019.06.036.

34. Schubert, U., Antón, L.C., Gibbs, J., Norbury, C.C., Yewdell, J.W., and Bennink, J.R. (2000). Rapid degradation of a large fraction of newly synthesized proteins by proteasomes. Nature 404, 770–774. 10.1038/35008096.

35. Kempf, B.J., and Barton, D.J. (2008). Poliovirus 2A Pro Increases Viral mRNA and Polysome Stability Coordinately in Time with Cleavage of eIF4G. J. Virol. 82, 5847–5859. 10.1128/JVI.01514-07.

36. Dassi, E., Zuccotti, P., Leo, S., Provenzani, A., Assfalg, M., D’Onofrio, M., Riva, P., and Quattrone, A. (2013). Hyper conserved elements in vertebrate mRNA 3′-UTRs reveal a translational network of RNA-binding proteins controlled by HuR. Nucleic Acids Res. 41, 3201–3216. 10.1093/nar/gkt017.

37. George, B., Dave, P., Rani, P., Behera, P., and Das, S. (2021). Cellular Protein HuR Regulates the Switching of Genomic RNA Templates for Differential Functions during the Coxsackievirus B3 Life Cycle. J. Virol. 95. 10.1128/JVI.00915-21.

38. Dever, T.E., Dinman, J.D., and Green, R. (2018). Translation Elongation and Recoding in Eukaryotes. Cold Spring Harb. Perspect. Biol. 10, a032649. 10.1101/cshperspect.a032649.

39. Roux, P.P., Shahbazian, D., Vu, H., Holz, M.K., Cohen, M.S., Taunton, J., Sonenberg, N., and Blenis, J. (2007). RAS/ERK Signaling Promotes Site-specific Ribosomal Protein S6 Phosphorylation via RSK and Stimulates Cap-dependent Translation. J. Biol. Chem. 282, 14056–14064. 10.1074/jbc.M700906200.

40. Pochopien, A.A., Beckert, B., Kasvandik, S., Berninghausen, O., Beckmann, R., Tenson, T., and Wilson, D.N. (2021). Structure of Gcn1 bound to stalled and colliding 80S ribosomes. Proc. Natl. Acad. Sci. 118. 10.1073/pnas.2022756118.

41. Blyn, L.B., Towner, J.S., Semler, B.L., and Ehrenfeld, E. (1997). Requirement of poly(rC) binding protein 2 for translation of poliovirus RNA. J. Virol. 71, 6243–6246. 10.1128/jvi.71.8.6243-6246.1997.

42. Han, S., Wang, X., Guan, J., Wu, J., Zhang, Y., Li, P., Liu, Z., Abdullah, S.W., Zhang, Z., Jin, Y., et al. (2021). Nucleolin Promotes IRES-Driven Translation of Foot-and-Mouth Disease Virus by Supporting the Assembly of Translation Initiation Complexes. J. Virol. 95. 10.1128/JVI.00238-21.

43. Levengood, J.D., Tolbert, M., Li, M.-L., and Tolbert, B.S. (2013). High-affinity interaction of hnRNP A1 with conserved RNA structural elements is required for translation and replication of enterovirus 71. RNA Biol. 10, 1136–1145. 10.4161/rna.25107.

44. Dobrikov, M.I., Dobrikova, E.Y., McKay, Z.P., Kastan, J.P., Brown, M.C., and Gromeier, M. (2022). PKR Binds Enterovirus IRESs, Displaces Host Translation Factors, and Impairs Viral Translation to Enable Innate Antiviral Signaling. MBio 13. 10.1128/mbio.00854-22.

45. Bedard, K.M., Daijogo, S., and Semler, B.L. (2007). A nucleo-cytoplasmic SR protein functions in viral IRES-mediated translation initiation. EMBO J. 26, 459–467. 10.1038/sj.emboj.7601494.

46. Chen, J., Zhang, R., Lan, J., Lin, S., Li, P., Gao, J., Wang, Y., Xie, Z.-J., Li, F.-C., and Jiang, S.-J. (2019). IGF2BP1 Significantly Enhances Translation Efficiency of Duck Hepatitis A Virus Type 1 without Affecting Viral Replication. Biomolecules 9, 594. 10.3390/biom9100594.

47. Hunt, S.L., Hsuan, J.J., Totty, N., and Jackson, R.J. (1999). unr, a cellular cytoplasmic RNA-binding protein with five cold-shock domains, is required for internal initiation of translation of human rhinovirus RNA. Genes Dev. 13, 437–448. 10.1101/gad.13.4.437.

48. Cathcart, A.L., Rozovics, J.M., and Semler, B.L. (2013). Cellular mRNA Decay Protein AUF1 Negatively Regulates Enterovirus and Human Rhinovirus Infections. J. Virol. 87, 10423–10434. 10.1128/JVI.01049-13.

49. Merrill, M.K., and Gromeier, M. (2006). The Double-Stranded RNA Binding Protein 76:NF45 Heterodimer Inhibits Translation Initiation at the Rhinovirus Type 2 Internal Ribosome Entry Site. J. Virol. 80, 6936–6942. 10.1128/JVI.00243-06.

50. López-Ulloa, B., Fuentes, Y., Pizarro-Ortega, M.S., and López-Lastra, M. (2022). RNA-Binding Proteins as Regulators of Internal Initiation of Viral mRNA Translation. Viruses 14, 188. 10.3390/v14020188.

51. Martínez-Salas, E., Lozano, G., Fernandez-Chamorro, J., Francisco-Velilla, R., Galan, A., and Diaz, R. (2013). RNA-binding proteins impacting on internal initiation of translation. Int. J. Mol. Sci. 14, 21705–21726. 10.3390/ijms141121705.

52. Komar, A.A., and Hatzoglou, M. (2011). Cellular IRES-mediated translation. Cell Cycle 10, 229–240. 10.4161/cc.10.2.14472.

53. Solomon, O., Oren, S., Safran, M., Deshet-Unger, N., Akiva, P., Jacob-Hirsch, J., Cesarkas, K., Kabesa, R., Amariglio, N., Unger, R., et al. (2013). Global regulation of alternative splicing by adenosine deaminase acting on RNA (ADAR). RNA 19, 591–604. 10.1261/rna.038042.112.

54. Quin, J., Sedmík, J., Vukić, D., Khan, A., Keegan, L.P., and O’Connell, M.A. (2021). ADAR RNA Modifications, the Epitranscriptome and Innate Immunity. Trends Biochem. Sci. 46, 758–771. 10.1016/j.tibs.2021.02.002.

55. Zhu, T., Niu, G., Zhang, Y., Chen, M., Li, C.-Y., Hao, L., and Zhang, Z. (2023). Host-mediated RNA editing in viruses. Biol. Direct 18, 12. 10.1186/s13062-023-00366-w.

56. Pfaller, C.K., George, C.X., and Samuel, C.E. (2021). Adenosine Deaminases Acting on RNA (ADARs) and Viral Infections. Annu. Rev. Virol. 8, 239–264. 10.1146/annurev-virology-091919-065320.

57. Burrill, C.P., Westesson, O., Schulte, M.B., Strings, V.R., Segal, M., and Andino, R. (2013). Global RNA Structure Analysis of Poliovirus Identifies a Conserved RNA Structure Involved in Viral Replication and Infectivity. J. Virol. 87, 11670–11683. 10.1128/JVI.01560-13.

58. Peng, Z., Cheng, Y., Tan, B.C.-M., Kang, L., Tian, Z., Zhu, Y., Zhang, W., Liang, Y., Hu, X., Tan, X., et al. (2012). Comprehensive analysis of RNA-Seq data reveals extensive RNA editing in a human transcriptome. Nat. Biotechnol. 30, 253–260. 10.1038/nbt.2122.

59. Balvay, L., Rifo, R.S., Ricci, E.P., Decimo, D., and Ohlmann, T. (2009). Structural and functional diversity of viral IRESes. Biochim. Biophys. Acta - Gene Regul. Mech. 1789, 542–557. 10.1016/j.bbagrm.2009.07.005.

60. Kozisek, T., Samuelson, L., Hamann, A., and Pannier, A.K. (2023). Systematic comparison of nonviral gene delivery strategies for efficient co-expression of two transgenes in human mesenchymal stem cells. J. Biol. Eng. 17, 76. 10.1186/s13036-023-00394-0.

61. Sommer, B., Köhler, M., Sprengel, R., and Seeburg, P.H. (1991). RNA editing in brain controls a determinant of ion flow in glutamate-gated channels. Cell 67, 11–19. 10.1016/0092-8674(91)90568-J.

62. Aviner, R., Geiger, T., and Elroy-Stein, O. (2014). Genome-wide identification and quantification of protein synthesis in cultured cells and whole tissues by puromycin-associated nascent chain proteomics (PUNCH-P). Nat. Protoc. 9, 751–760. 10.1038/nprot.2014.051.

63. Xiao, Y., Dolan, P.T., Goldstein, E.F., Li, M., Farkov, M., Brodsky, L., and Andino, R. (2017). Poliovirus intrahost evolution is required to overcome tissue-specific innate immune responses. Nat. Commun. 8, 375. 10.1038/s41467-017-00354-5.

64. Zhang, K., Wang, S., Chen, T., Tu, Z., Huang, X., Zang, G., Wu, C., Fan, X., Liu, J., Tian, Y., et al. (2022). ADAR1p110 promotes Enterovirus D68 replication through its deaminase domain and inhibition of PKR pathway. Virol. J. 19, 222. 10.1186/s12985-022-01952-6.

65. Zhou, S., Yang, C., Zhao, F., Huang, Y., Lin, Y., Huang, C., Ma, X., Du, J., Wang, Y., Long, G., et al. (2019). Double-stranded RNA deaminase ADAR1 promotes the Zika virus replication by inhibiting the activation of protein kinase PKR. J. Biol. Chem. 294, 18168–18180. 10.1074/jbc.RA119.009113.

66. Okonski, K.M., and Samuel, C.E. (2013). Stress granule formation induced by measles virus is protein kinase PKR dependent and impaired by RNA adenosine deaminase ADAR1. J. Virol. 87, 756–766. 10.1128/JVI.02270-12.

67. Zahn, R.C., Schelp, I., Utermöhlen, O., and von Laer, D. (2007). A-to-G Hypermutation in the Genome of Lymphocytic Choriomeningitis Virus. J. Virol. 81, 457–464. 10.1128/JVI.00067-06.

68. Piontkivska, H., Wales-McGrath, B., Miyamoto, M., and Wayne, M.L. (2021). ADAR Editing in Viruses: An Evolutionary Force to Reckon with. Genome Biol. Evol. 13. 10.1093/gbe/evab240.

69. Freund, E.C., Sapiro, A.L., Li, Q., Linder, S., Moresco, J.J., Yates, J.R., and Li, J.B. (2020). Unbiased Identification of trans Regulators of ADAR and A-to-I RNA Editing. Cell Rep. 31, 107656. 10.1016/j.celrep.2020.107656.

70. Khadka, S., Williams, C.G., Sweeney-Gibbons, J., and Basler, C.F. (2021). Marburg and Ebola Virus mRNA 3′ Untranslated Regions Contain Negative Regulators of Translation That Are Modulated by ADAR1 Editing. J. Virol. 95. 10.1128/JVI.00652-21.

71. Yin, C., Zhao, H., Xia, X., Pan, Z., Li, D., and Zhang, L. (2024). Picornavirus 2C proteins: structure-function relationships and interactions with host factors. Front. Cell. Infect. Microbiol. 14, 1347615. 10.3389/fcimb.2024.1347615.

72. Shimizu, H., Agoh, M., Agoh, Y., Yoshida, H., Yoshii, K., Yoneyama, T., Hagiwara, A., and Miyamura, T. (2000). Mutations in the 2C Region of Poliovirus Responsible for Altered Sensitivity to Benzimidazole Derivatives. J. Virol. 74, 4146–4154. 10.1128/JVI.74.9.4146-4154.2000.

73. Li, D., and Zhang, L. (2022). Structure Prediction and Potential Inhibitors Docking of Enterovirus 2C Proteins. Front. Microbiol. 13. 10.3389/fmicb.2022.856574.

74. Fitzsimmons, W.J., Woods, R.J., McCrone, J.T., Woodman, A., Arnold, J.J., Yennawar, M., Evans, R., Cameron, C.E., and Lauring, A.S. (2018). A speed-fidelity trade-off determines the mutation rate and virulence of an RNA virus. PLoS Biol. 16, e2006459. 10.1371/journal.pbio.2006459.

75. Wright, A.L., Konen, L.M., Mockett, B.G., Morris, G.P., Singh, A., Burbano, L.E., Milham, L., Hoang, M., Zinn, R., Chesworth, R., et al. (2023). The Q/R editing site of AMPA receptor GluA2 subunit acts as an epigenetic switch regulating dendritic spines, neurodegeneration and cognitive deficits in Alzheimer’s disease. Mol. Neurodegener. 18, 65. 10.1186/s13024-023-00632-5.

76. DeJesus, N., Franco, D., Paul, A., Wimmer, E., and Cello, J. (2005). Mutation of a Single Conserved Nucleotide between the Cloverleaf and Internal Ribosome Entry Site Attenuates Poliovirus Neurovirulence. J. Virol. 79, 14235–14243. 10.1128/JVI.79.22.14235-14243.2005.

77. Evans, D.M.A., Dunn, G., Minor, P.D., Schild, G.C., Cann, A.J., Stanway, G., Almond, J.W., Currey, K., and Maizel, J. V. (1985). Increased neurovirulence associated with a single nucleotide change in a noncoding region of the Sabin type 3 poliovaccine genome. Nature 314, 548–550. 10.1038/314548a0.

78. Liu, Y., Ma, T., Liu, J., Zhao, X., Cheng, Z., Guo, H., Xu, R., and Wang, S. (2015). Circulating type 1 vaccine-derived poliovirus may evolve under the pressure of adenosine deaminases acting on RNA. J. Matern. Neonatal Med. 28, 2096–2099. 10.3109/14767058.2014.979147.

79. Hideyama, T., Yamashita, T., Aizawa, H., Tsuji, S., Kakita, A., Takahashi, H., and Kwak, S. (2012). Profound downregulation of the RNA editing enzyme ADAR2 in ALS spinal motor neurons. Neurobiol. Dis. 45, 1121–1128. 10.1016/j.nbd.2011.12.033.

80. Hur, S. (2019). Double-Stranded RNA Sensors and Modulators in Innate Immunity. Annu. Rev. Immunol. 37, 349–375. 10.1146/annurev-immunol-042718-041356.

81. Perez-Riverol, Y., Csordas, A., Bai, J., Bernal-Llinares, M., Hewapathirana, S., Kundu, D.J., Inuganti, A., Griss, J., Mayer, G., Eisenacher, M., et al. (2019). The PRIDE database and related tools and resources in 2019: improving support for quantification data. Nucleic Acids Res. 47, D442–D450. 10.1093/nar/gky1106.

82. Gamarnik, A. V, and Andino, R. (1998). Switch from translation to RNA replication in a positive-stranded RNA virus. Genes Dev. 12, 2293–2304. 10.1101/gad.12.15.2293.

83. Burrill, C.P., Strings, V.R., and Andino, R. (2013). Poliovirus: generation, quantification, propagation, purification, and storage. Curr. Protoc. Microbiol. Chapter 15, 15H.1.1–15H.1.27. 10.1002/9780471729259.mc15h01s29.

84. Xiao, Y., Dolan, P.T., Goldstein, E.F., Li, M., Farkov, M., Brodsky, L., and Andino, R. (2017). Poliovirus intrahost evolution is required to overcome tissue-specific innate immune responses. Nat. Commun. 8, 375. 10.1038/s41467-017-00354-5.

85. Ingolia, N.T., Brar, G.A., Rouskin, S., McGeachy, A.M., and Weissman, J.S. (2012). The ribosome profiling strategy for monitoring translation in vivo by deep sequencing of ribosome-protected mRNA fragments. Nat. Protoc. 7, 1534–1550. 10.1038/nprot.2012.086.

86. Wilm, A., Aw, P.P.K., Bertrand, D., Yeo, G.H.T., Ong, S.H., Wong, C.H., Khor, C.C., Petric, R., Hibberd, M.L., and Nagarajan, N. (2012). LoFreq: a sequence-quality aware, ultra-sensitive variant caller for uncovering cell-population heterogeneity from high-throughput sequencing datasets. Nucleic Acids Res. 40, 11189–11201. 10.1093/nar/gks918.

87. Rappsilber, J., Mann, M., and Ishihama, Y. (2007). Protocol for micro-purification, enrichment, pre-fractionation and storage of peptides for proteomics using StageTips. Nat. Protoc. 2, 1896–1906. 10.1038/nprot.2007.261.

88. Cox, J., and Mann, M. (2008). MaxQuant enables high peptide identification rates, individualized p.p.b.-range mass accuracies and proteome-wide protein quantification. Nat. Biotechnol. 26, 1367–1372. 10.1038/nbt.1511.

89. Cox, J., Neuhauser, N., Michalski, A., Scheltema, R.A., Olsen, J. V, and Mann, M. (2011). Andromeda: a peptide search engine integrated into the MaxQuant environment. J. Proteome Res. 10, 1794–1805. 10.1021/pr101065j.

90. Tusher, V.G., Tibshirani, R., and Chu, G. (2001). Significance analysis of microarrays applied to the ionizing radiation response. Proc. Natl. Acad. Sci. 98, 5116–5121. 10.1073/pnas.091062498.

91. Meng, E.C., Goddard, T.D., Pettersen, E.F., Couch, G.S., Pearson, Z.J., Morris, J.H., and Ferrin, T.E. (2023). <scp>UCSF ChimeraX</scp> : Tools for structure building and analysis. Protein Sci. 32. 10.1002/pro.4792.

